# Linking Expression and Function of *Drosophila* Type-I TGF-β Receptor Baboon Isoforms: Multiple Roles of BaboA Isoform in Shaping of the Adult Central Nervous System

**DOI:** 10.1101/2025.01.17.633552

**Authors:** Gyunghee G. Lee, Aidan J. Peterson, Myung-Jun Kim, MaryJane Shimell, Michael B. O’Connor, Jae H. Park

## Abstract

Evolutionarily conserved transforming growth factor β (TGF-β) signaling is used in both vertebrates and invertebrates to regulate a variety of developmental and cellular processes. The *baboon* (*babo*) gene encoding a *Drosophila* type-I TGF-β receptor produces three isoforms via alternative splicing: BaboA, BaboB, and BaboC. In this study, we generated three fly lines, each carrying an isoform-specific GFP tag, and another line with a GFP conjugated at the C-terminus common to all isoforms. Using these lines, we assessed (1) whether the tagged proteins function properly in rescue assays and (2) how the isoform expression is regulated in various tissues including the central nervous system (CNS). A *Gal4* knock-in line in the *babo* locus was also characterized for reporter expression, mutant phenotypes, and isoform-specific knockdown phenotypes. We found that the C-terminal tag does not interrupt the subcellular targeting and functions of the tagged isoforms, but the internal isoform tags do so in a cell- and isoform-specific fashion. Nevertheless, our results demonstrated that these tags faithfully reflect endogenous expression of individual isoforms. Certain cell types express single or multiple isoforms at different levels, suggesting that alternative splicing could determine the isoform types and their levels depending on cell (or tissue) type. The larval CNS displays distinct patterns of two isoforms, BaboA and BaboC. BaboC is mostly expressed in neural cells originating during embryogenesis, while BaboA is broadly expressed in neural cells produced from both embryonic and postembryonic stages. Assays of both isoform-specific mutants and cell-specific knockdown of individual isoforms revealed broad roles played by BaboA in postembryonic neurogenesis and differentiation of precursor neurons, remodeling processes of persisting larval neurons, and metamorphic CNS reorganization, which are essential for establishing of the adult CNS. Taken together, this study demonstrates that the GFP-tagged lines permit visualization of endogenous expression of individual isoforms, which further provides clues about cell- and stage-specific functions played by each isoform.

## Introduction

The TGF-β signaling is evolutionarily conserved signal transduction module that works in a contextual manner to mediate diverse cellular processes including growth, differentiation, metabolism, programmed cell death (PCD), and morphogenesis of both vertebrates and invertebrates [1–3]. As in mammals, *Drosophila* TGF-β signaling is sorted into two branches, Activin and Bone morphogenetic protein (BMP) signalings, each utilizing specific type-I receptors activated by distinct ligands, and different downstream R-Smad transcriptional transducers, dSmad2 and Mad, respectively [2, 4, 5]. In the Activin branch, three ligands, Activin-β (Actβ), Dawdle (Daw) and Myoglianin (Myo), are suggested to signal through the type-I receptor Babo. Notably the *babo* gene produces three isoforms, BaboA, BaboB, and BaboC, which differ only in ligand binding domains. A growing body of studies using genetic, transgenic, and cell-based analyses support the following ligand/Babo isoform pairings: Myo/BaboA, Actβ/BaboB, and Daw/BaboC [2, 6–10].

Prior genetic studies have suggested that the Babo isoforms act individually and cooperatively in certain aspects of larval brain development [10]. However, how expression of the individual isoforms is regulated has not been reported in this organ. All Activin branch ligands are expressed in the developing CNS at late 3^rd^ instar larval stage. Expression of Actβ is seen in several neuronal subtypes including mushroom body, peptidergic and motor neurons [11, 12]. Both Daw and Myo are expressed exclusively in glial cells within the CNS; Daw in perineurial glia [11, 13], and Myo in cortex and astrocyte-like glia subtypes [7]. To fully understand the biological roles of the versatile TGF-β signaling, it is crucial to know the target tissues/cells expressing their cognate receptors. Considering expression of the ligands in the CNS, one would expect their cognate receptors to be found within this organ as well. For instance, Daw is shown to act on the insulin producing cells in the brain of feeding 3^rd^ instar larvae to regulate release of the *Drosophila* insulin-like proteins (Dilps) [8]. However, it is not resolved whether Daw acts by Daw-BaboC transduction in these cells. Cell- and stage-specific knockdown (KD) assays revealed an autonomous role of BaboA in the NE-to-NB conversion event [10]. Similar approaches have shown Myo-BaboA action in a few metamorphosis-associated cellular events including remodeling processes of the mushroom body (MB) γ neurons [7] and developmental PCD of the vCrz neurons that occurs during early phase of metamorphosis [9]. Nevertheless, these limited studies are insufficient to fully comprehend the CNS functions of individual Babo isoforms.

The cellular locations of the Babo isoforms provide invaluable clues about isoform functions and how cellular networks of each isoform-mediated signaling are formed. At the whole tissue level, RT-PCR has revealed that BaboC is the predominant form in the fat body, while BaboA is the major one in imaginal discs and brains [14]. However, this analysis is insufficient to determine if there are small subpopulations of cells within the tissue that express different isoforms at low levels. One such tissue type is the CNS that consists of immensely heterogenous cell types in origin, structure, and function.

Previously, a knock-in fly line carrying a human influenza hemagglutinin (HA) tag at the C-terminus of Babo was employed to detect endogenously produced Babo::HA by using anti-HA, revealing the protein primarily enriched in neurites of the MB and the optic lobe (OL) [15]. However, this strategy proves a difficulty to further resolve other Babo-expressing cell types in detail due to very low levels of Babo proteins. Furthermore, the common C-terminal tag does not permit the identification of isoforms expressed. Here, we generated GFP-tagged reporter transgenic lines of *babo*. The GFP-tag constructs were engineered to detect endogenous production of each isoform (isoform-specific) as well all Babo isoforms together (pan-Babo) [16, 17]. Using these lines, we explored how *babo* isoform expression is regulated in various tissues including the CNS and the ring gland. These observations, together with assays of both isoform-specific null mutations and cell-specific KD of the isoforms revealed that alternative splicing determines isoform types and their differential levels in a cell-specific manner and that BaboA plays a broad role to reconstruct, in both structure and function, the larval CNS into that of an adult one.

## Materials and Methods

### Fly strains

Flies were raised on standard cornmeal-yeast-agar food at 25°C unless otherwise specified. For transgenic manipulations, we used *scarecrow* (*scro*)^ΔE2-*Gal4*^ (*scro-Gal4*) [18], *e22c*-*Gal4* [19], and *babo^CR00274-TG4.1^* (*babo^Gal4^*, BL# 83164). To visualize *Gal4* activities, three reporters, *UAS-mCD8GFP* (BL#5136), *UAS-stinger* (a.k.a. *UAS-nGFP* or *UAS-GFPnls*, BL#4775), and *UAS-redstinger* (a.k.a. *UAS-nRFP*, BL#8547) were used. Isoform-specific deletion mutations, *baboA-indel*, *baboB-indel*, and *baboC-indel*, were as described [10]. For isoform-specific KD assays, we used *UAS-baboA-miRNA*, *UAS-baboB-miRNA*, *UAS-baboC-miRNA* lines inserted at *attp16* (mainly used and referred to as #1) [7] or *VK37* site (referred to as #2). A *trans* combination of *babo^Fd4^* and *babo^Df^* (*babo^Fd4/Df^*) was used as the *babo*-null mutation and *w^1118^* as a genetic control [10]. BL indicates the stocks obtained from the Bloomington Drosophila Stock Center (https://flystocks.bio.indiana.edu/).

### Generation of GFP-tagged transgenic lines of *babo*

All forms of *babo* were tagged using recombineering technology [20]. The starting BAC clone (CH322-78I5) was obtained from BacPac resources (https://bacpacresources.org/home.htm). For a pBac{pan-*babo*-*GFP*} construct, GFP was appended to the C-terminus in-frame using recombineering as described [16, 17]. For a comparison, we obtained and tested similarly generated line (*babo^fTRG00444.sfGFP-TVPTBF^*, id. 318433) from Vienna Drosophila Resource Center [21]. Isoforms A, B, and C were tagged in their respective fourth exons between the Cys box (CCKSDFCN) and the end of the exon, before the splice donor site, to generate in-frame isoform-specific GFP tags. Specifically, the GFP of BaboA was inserted between Ile205 and Met206, 21 bp from the end of exon 4; the GFP of BaboB between Ile230 and Ser231, 9 bp from the end of exon 4; the GFP of BaboC between Ser202 and Gly203, 10 bp from the end of exon 4. The GFP template plasmid was PL-452 N-EGFP (AddGene #19173). Amplifying primers consisted of 50 bases of *babo* sequence fused to *N EGFP for* and 50 bases of *babo* sequence fused to *N term PL452 rev*. Primers are listed in Supplemental Table 1. The PCR selection cassettes were recombined onto CH322-78I5 in cells. Arabinose induction of *loxP* excision led to an in-frame *loxP* GFP cassette in the *babo* fourth exon. Transgenic founders were recovered after injection into the *attP2* (3^rd^ chromosome) docking site stock (BestGene).

### Immunohistochemistry

Immunohistochemistry for the CNS and the ring gland tissues followed the procedures as described [22], except for 2-h fixation with shaking at room temperature (RT). For immunostaining of body wall tissues, actively wandering larvae were rinsed in dH_2_O, dissected in Ca^2+^-free 1X PBS, then fixed in 3.8 % Paraformaldehyde solution (Electron Microscopy Sciences) for 40 min at RT [23]. After washing in 1X PBS and permeabilization in 1X PBT (0.5% BSA+0.2% Triton X-100 in 1X PBS), the tissues were incubated with primary antibodies. The following primary antibodies were used: rabbit-anti-GFP (1:500) [24], mouse-anti-24B10 (1:100, Developmental Studies Hybridoma, Bank, DSHB), mouse-anti-Dlg (1:100, DSHB 4F3), rat-anti-Mira (1:100, Abcam), mouse-anti-EcR-B1 (1:20, DSHB AD4.4), rat-anti-Elav (1:100, DSHB), mouse-anti-Repo (1:100 DSHB,), anti-Prospero (1:100, DSHB), rabbit-anti-PDF (1: 3000) [22], rabbit-anti-Crz [1: 3000] [25], rat-anti-Dilp2 (1/500) [26], and mouse-anti-FasIII (1:100, DSHB, 7G10). Rabbit polyclonal anti-NPF (1:600, antigenic region: YKFLQDLDTYYGDR) and anti-DSK (1:600, antigenic region: IELDLLMDNDDERTK) were generated for this study (Genemed Synthesis). Secondary antibodies conjugated with Alex Fluor 594 (1:200; Jackson ImmunoResearch Lab), Alex Fluor 555 (1:200, Molecular Probes), or Alex Fluor 488 (1:200, Molecular Probes) were used. DAPI staining was done by treating tissues for 1 h after the final 4^th^ wash in PBS containing 0.02% Triton X-100. The processed CNS tissues were mounted with a shielding medium (2% n-propyl galate in 80% glycerol, 20% of 1X PBS). Confocal images were obtained with a Zeiss LSM710 or Leica SP8 confocal microscope. Epi-fluorescent and bright-field images were taken with an Olympus BX61 microscope. Fly body images were taken with a Nikon stereo microscope SMZ645 using Seba ViewLife program. Figures were generated using ImageJ (NIH) and Microsoft PowerPoint.

### Immunoblot analysis

Body wall tissues of actively wandering larvae were homogenized in RIPA buffer (Sigma, #R0278) supplemented with a cocktail of protease inhibitors (Complete mini, Roche) and incubated at 4°C for 40 min with agitation. After centrifugation, supernatants were transferred into new tubes, mixed with 3X loading buffer and denatured for 5 min at 95°C. Equal volumes from each sample were run on 4-12% Bis-Tris gels (Novex, #NP0322BOX) and proteins were transferred to PVDF membranes (Millipore, #IPFL00010). The membranes were then blocked with Casein-containing buffer (Bio-Rad, #1610783) and incubated with primary antibodies at 4°C overnight. The primary antibodies were rabbit anti-pSmad2 (1:500, Cell signaling Technology, #3108) and anti-β-Tubulin (1:1000, DSHB, E7). The immunoreactive bands were detected using HRP-conjugated secondary antibodies (anti-rabbit and anti-mouse IgGs, 1:10,000, Cell signaling Technology), developed using Pierce ECL Western Blotting Substrate (Thermo Scientific, #32209), and chemiluminescence detected after exposure to X-ray film (Genesee Scientific). Relative band intensities were quantified using Image J (NIH) software. The quantification graph showing relative protein levels is presented beneath the representative immunoblot and data are mean ± sem from three independent samplings.

### Measuring areas of brain regions

For accurate developmental staging, larvae were raised on a special apple juice medium as descried [8, 10]. Larval brains were dissected at 120 h after egg laying (AEL), processed for anti-Dlg and DAPI staining, and mounted with dorsal side up between two 1-mm thick 18×18 mm coverslips as spacers on a microscope slide, covered with another coverslip on top, and sealed with clear nail polish. A mid z-section (2 μm thick) image was taken from one brain lobe per sample. Regional areas were manually measured on each z-section image using the Image J program.

### Statistics

Statistical analyses were performed using GraphPad Prism (v9.0) or Instat (v2.0). Ordinary one-way ANOVA followed by multiple comparisons was used for more than three groups and unpaired student t-test for comparison between two groups. Asterisks denote statistical significance with the following p-vales: *p<0.05, ** p<0.01, ***p<0.001, ****p<0.0001, and ns for not significant (p>0.05). Graphs were generated by using either GraphPad Prism or Microsoft Excel. All bars in the graphs show mean ± sd (standard deviation) unless otherwise indicated.

## Results

### GFP-tags to detect pan-Babo or individual isoforms

The *babo* gene contains three isoform-specific 4^th^ exons (e4-A/B/C), each of which encodes a distinct extracellular ligand binding domain (Fig 1A and 1B). Therefore, we predict alternative splicing as the key mechanism that determines differential expression patterns of the isoforms. To investigate how expression of the isoform is regulated, we generated reporter lines, each carrying a genomic *babo* fosmid clone modified by an in-frame insertion of GFP in the 4^th^ exon (Fig 1C, details in Materials and Methods). These lines are labeled as *baboA-GFP*, *baboB-GFP*, and *baboC-GFP*. To assess the sum expression of all isoforms, we also made an additional line containing a C-terminal GFP-tag; the line carrying this construct is hereafter referred to as pan-*babo-GFP* (Fig 1C). Homozygous stocks did not show noticeable defects, signifying that the additional *babo* locus is not detrimental to general development and adult performance.

**Fig 1.**
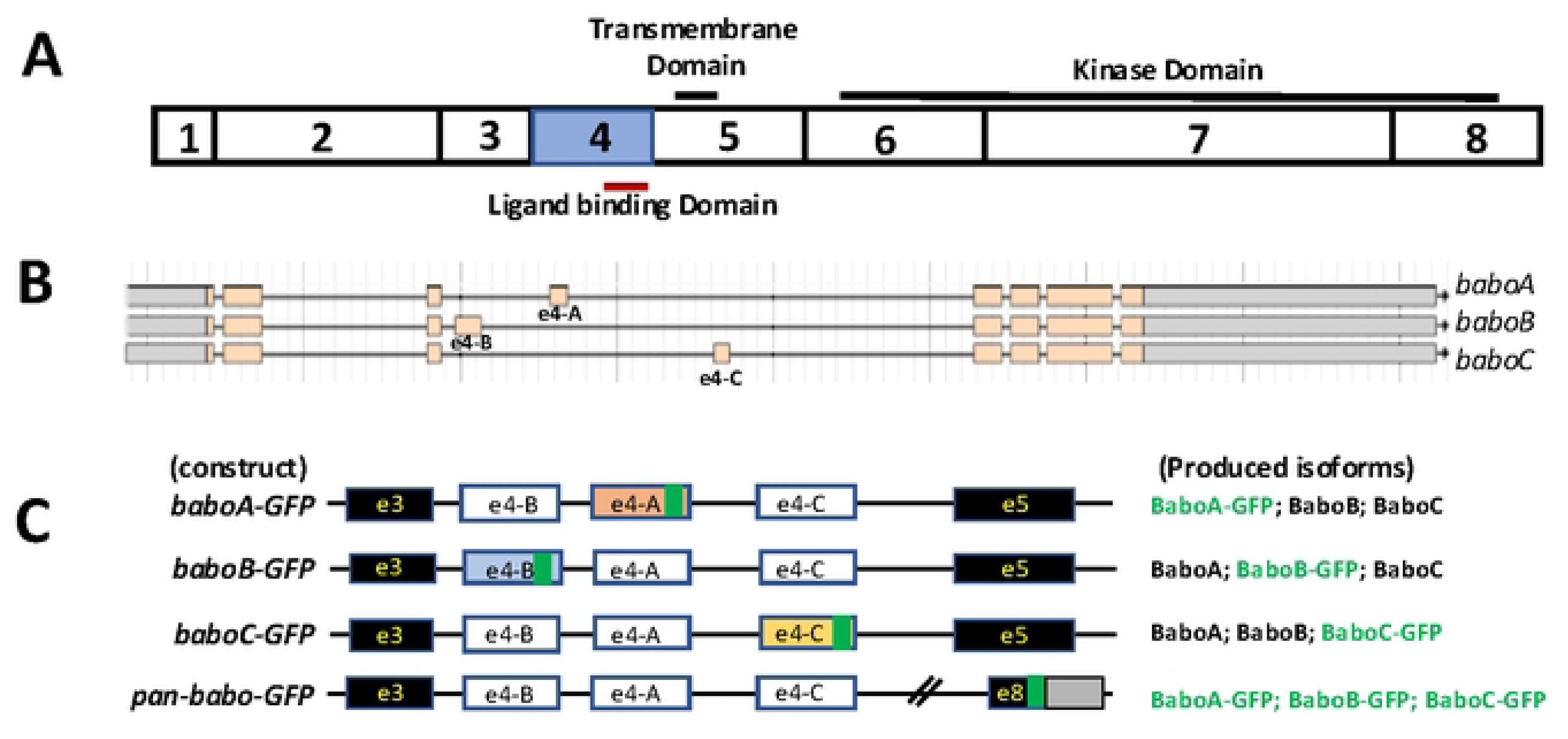
Generation of fly lines, each carrying a GFP-tag construct to label individual or all Babo isoforms. (A) Coding exons of *babo* gene. Three domains for its function as the receptor are indicated on the solid lines. (B) Genomic structure of the *babo* locus (boxes indicate exons) from the Flybase. Alternative 4^th^ exon (e4-A/B/C) produces three *babo* isoforms. (C) Schematics of the constructs showing in-frame GFP insertions (green box) within the isoform-specific 4^th^ exons (white) and at the C-terminus of the common 8^th^ exon (e8) that contains both coding (black) and non-coding (gray) regions. In *pan-babo-GFP*, all isoforms are expected to be C-terminally tagged, while isoform-specific constructs produce a respective GFP-tagged isoform (green) and two other non-tagged isoforms (black).

### Rescue assays

As an initial assessment of functional consequence of the in-frame GFP insertions, we tested the ability of the tagged constructs to provide *bab*o activity in genetic rescue experiments. Animals bearing a rescue construct in an otherwise *babo* loss-of-function context (*babo^Fd4/Df^*) were scored for survival to adulthood. As a negative control, we confirmed that *babo^Fd4/Df^* mutants mostly die during larval stages (Table 1). In contrast, one copy of the *pan-babo-GFP* provided significant rescue to adulthood, confirming that this construct drives sufficient expression of all GFP-tagged isoforms and that the tag does not interfere their receptor function (Table 1). We then compared the rescue strength of the tagged isoforms. The *baboA-* specific null (*A-indel*) mutants display severe defects during early phases of metamorphosis; only a small population manages to reach the pharate stage with morphological defects and dies without adult emergence, indicating that this isoform is vital for metamorphosis and eclosion [10]. The *baboA-GFP* construct, which is expected to produce GFP-tagged BaboA and non-tagged BaboB and BaboC (Fig 1C), phenocopies *baboA-indel* mutants. The simplest interpretation is that the GFP tag disrupts the function of the BaboA. The *baboB-GFP* construct, however, provides rescue comparable to *pan-babo-GFP*. Since the loss of BaboB function compromises viability at the pharate stage [10], this result suggests that the BaboB-GFP is less disruptive or does not affect the tagged isoform function. *baboC-GFP* displays no rescue on standard cornmeal food but enables some *babo* mutants to survive on low sugar food. This matches the sugar-sensitive behavior of *daw* or *baboC-indel* mutants [8, 10] and thus indicates that this construct behaves as only a loss-of-function for the C-isoform.

**Table 1:** Genetic rescue of *babo^Fd4/Df^* animals to ‘adulthood’ by one copy of various GFP-tagged constructs.

| Food | Construct: | N/A | Pan | A | B | C |
| --- | --- | --- | --- | --- | --- | --- |
| CMF | Rescue Index: | 0.0 | 0.76 | 0.0 | 0.84 | 0.0 |
|  | Rescue/Total: | 0/212 | 113/444 | 0/345 | 59/210 | 0/349 |
| 5Y5S | Rescue Index: | 0.0 | 0.81 | 0.0 | 0.67 | 0.08 |
|  | Rescue/Total: | 0/411 | 124/460 | 0/439 | 68/303 | 12/442 |

Larval and pupal phenotypes of the isoform-tag rescue animals provided further support for the above-mentioned results. The larval phenotypes of rescued animals are shown in Figure 2 in comparison to the *babo-null* mutant one. One copy of *pan-babo-GFP* restores normal size and shape of larvae and pupae and rescues the swollen anal pad phenotype (Fig 2A vs. 2B; 2F vs. 2G; 2K). Consistent with the lack of adult viability seen in *baboA-GFP* rescue assays, these animals fail to pupate properly and display defective puparia, supporting a loss-of-function of the tagged isoform (Fig 2C and 2H). The *baboB-GFP* animals form normal-sized larvae and pupae (Fig 2D and 2I), which is consistent with the interpretation that this GFP tag is not disruptive. *baboC-GFP* animals retain the distinctive anal pad phenotype but are otherwise normal-sized larvae and pupae (Fig 2E and 2J). Because the swollen anal pads are a hallmark of defective Daw-BaboC signaling [27], this further strengthens the conclusion that the GFP insertion blocks the function of BaboC. These data together support for the rescue by *baboB-GFP* and for isoform-specific defects in *baboA-GFP* and *baboC-GFP*. A combination of *baboA-GFP* and *baboC-GFP* constructs rescues the lethality and other defects (Table 2), further supporting that *baboA-GFP* line can provide functional BaboB and C, and *baboC-GFP* can do BaboA and B. Given that the mechanism of alternative splicing of *babo* transcripts is unknown, it cannot be predicted if the tagging exons will alter splicing and have unintended consequences. Thus, we were careful in designing the GFP tags upstream of the splice donor sites of the 4^th^ exon not to affect splicing and our rescue results indeed support this. For instance, the *baboB-GFP* line shows no apparent loss-of-function, which indicates that this isoform is functional with the GFP tag, and the lack of BaboA- or BaboC-associated phenotypes of this construct further implies that there is no significant disruption to the expression of the isoforms. Moreover, the full rescue by a combination of *baboA-GFP* and *baboC-GFP* confirm that non-tagged isoforms are properly expressed and remain functional. These results bolster our confidence that the tags should provide faithful spatial and temporal readouts of which cells normally produce the splice isoforms of *babo*.

**Fig 2.**
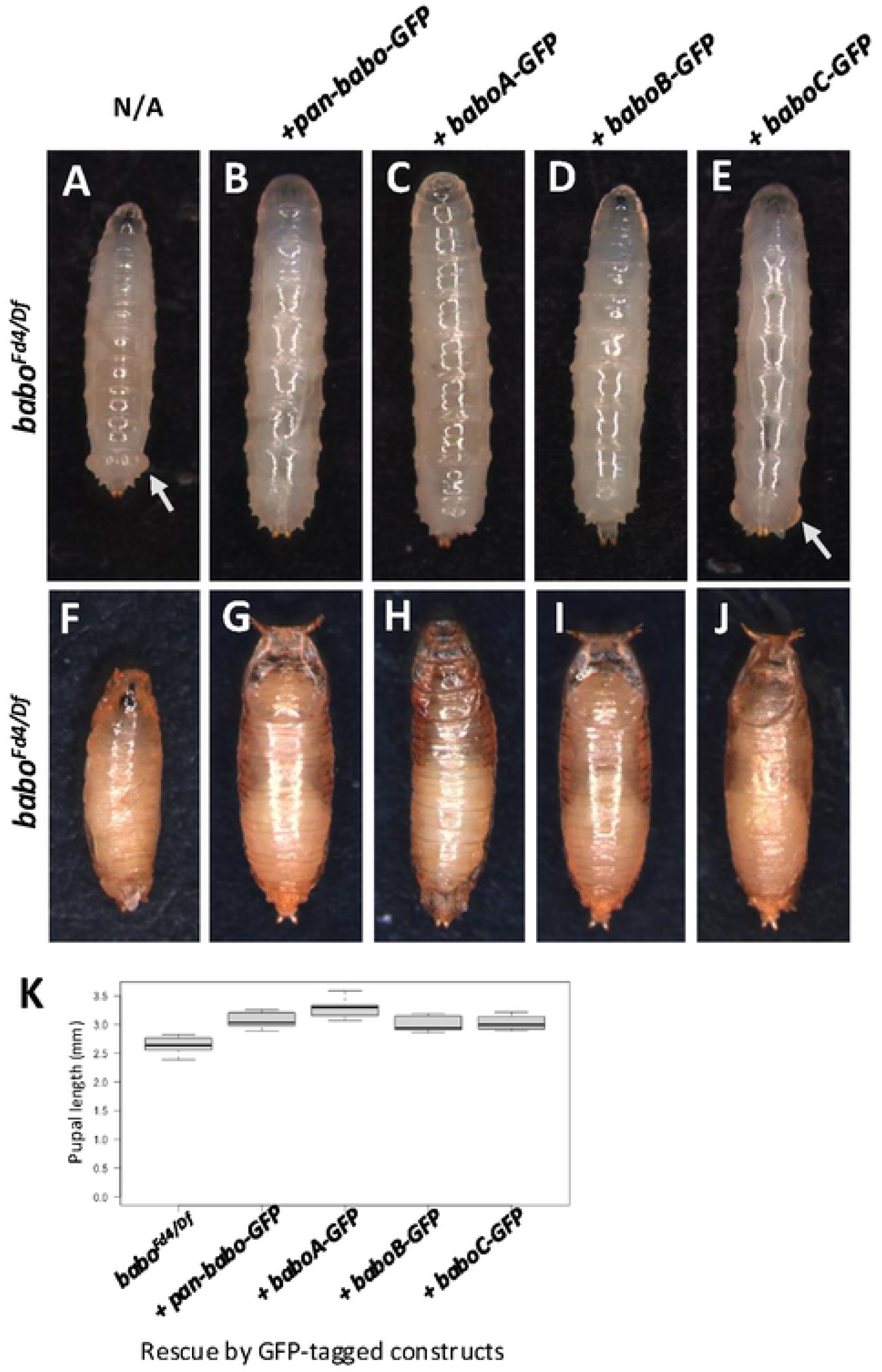
Larval and pupal morphology of rescued animals. (A-E) *babo^Fd4/Df^* mutant larvae rescued by indicated constructs. (A) The mutant alone is short with swollen anal pads (arrow). (B) Single copy of the *pan*-*babo-GFP* construct restores normal appearance. (C-E) Isoform-specific GFP tag constructs produce overtly normally sized larvae, but the *baboC-GFP* construct fails to rescue the swollen anal pad morphology (E, arrow). (F-J) Pupal appearance for the same genotype panel, imaged approximately four days after pupariation. The defective pupation of a *babo* mutant (F) is rescued by the *pan*-*babo-GFP* construct (G). Animals harboring *baboA-GFP* produced slightly elongated pupal cases with altered shape and failed to eclose (H). *baboB-* or *baboC-GFP* animals (I, J) are similar in appearance and size to those rescued by the positive control *pan*-*babo-GFP* construct. (K) Quantification of the pupal length for each genotype. The GFP-tagged constructs rescued the *babo* mutants except for *baboA-GFP* that is slightly longer than pan-*babo-GFP*, whereas *baboB-* and *baboC-GFP* are not statistically different from *pan*-*babo-GFP* animals.

**Table 2:** Genetic rescue of *babo^Fd4/Df^* animals to ‘adulthood’ by two copy of various GFP-tagged constructs.

| Construct: | A x A | C x C | A x C |
| --- | --- | --- | --- |
| Rescue Index: | 0.0 | 0.0 | 0.66 |
| Rescue/Total: | 0/621 | 0/1006 | 206/931 |

### pan-Babo and isoform-specific expression in the ring gland

Previous assays using prothoracic gland (PG)-specific KD and overexpression of individual key members of TGF-β signaling have suggested Babo functioning in this gland to modulate a stage-specific rise in ecdysone titer for triggering metamorphosis [28]. To gain histological evidence for this, we assessed *pan-babo-GFP* expression in the larval ring gland (RG), which consists of PG and two additional endocrine tissues, the corpora alata (CA), and the corpora cardiaca (CC). Individual endocrine cell types are easily discernible based on their relative positions and somatal sizes [29–32]. Since the GFP signal alone was too weak to visualize the tagged GFP even in the tissues bearing two copies of this construct, we employed polyclonal anti-GFP. The anti-GFP signal is undetectable in the PG of *w^1118^* but strong in the *pan-babo-GFP* one, supporting Babo expression in this gland as well as the specificity of anti-GFP (Fig 3A and 3B). In addition, we found GFP signals in the CA and CC, leading to a speculation that Babo might play certain roles in regulating the endocrine function of these tissues as well (Fig 3B).

**Fig 3.**
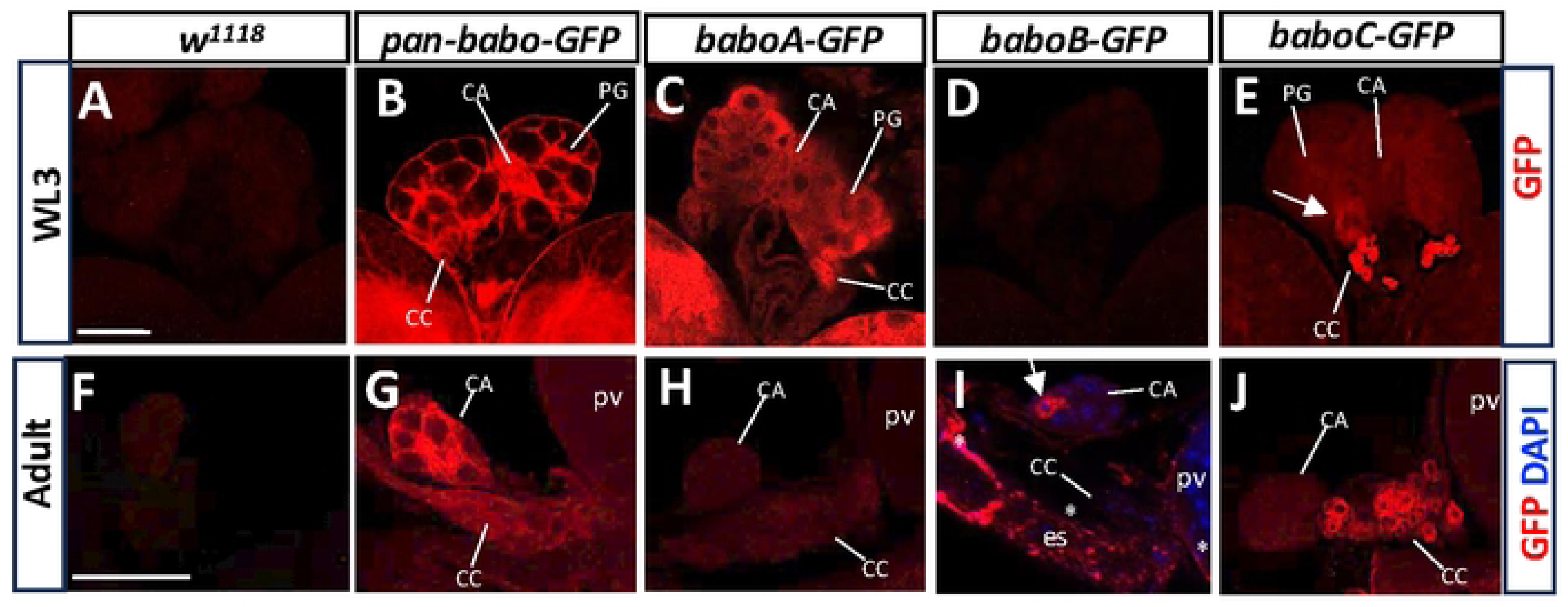
Expression of GFP-tagged Babo proteins by using anti-GFP in the ring gland. *w^1118^* was used as control. (A-E) More than 14 samples per genotype were examined for WL3 stage with two separate trials, and (F-J) 6-7 samples for adult stage. Prothoracic gland (PG), CA (corpora alata), and CC (corpora cardiaca) are indicated by lines. An arrow in (E) indicates PG cells with slightly elevated *baboC-GFP* expression levels. Asterisks in (I) indicate *baboB-GFP* on the cellular membrane of esophagus (es) and proventriculus (pv) cells, and an arrow points to cytoplasmic BaboB-GFP in a CA cell. Scale bars in A and F, 50 μm.

To resolve which isoforms are represented by the pan-Babo-GFP staining, we examined the larval RG of the isoform-tag lines. Intense expression of BaboA-GFP is seen in the three endocrine types (Fig 3C), whereas BaboB-GFP is absent in any of the larval RG tissues (Fig 3D). BaboC-GFP is also expressed in all RG tissues, although its expression levels are variable among tissue types: weak in the PG and CA, but strong in the CC (Fig 3E). Especially, about 50% of the specimens showed a few PG cells with noticeably elevated BaboC-GFP expression in a stochastic manner (indicated by an arrow in Fig 3E). The nature of such variable BaboC-GFP expression among PG cells is currently unknown but heterogenous expression of the ecdysone biosynthetic enzymes in the PG has been noted previously [28].

Larval PG cells undergo PCD during metamorphosis, while the CA and CC persist into adulthood but relocate to near the end of the esophagus (es) region, close to the proventriculus (pv) where the crop duct branches off [29, 32]. We sought further to see if the isoform expression patterns remain similar in the adult ones. Strong pan-Babo-GFP expression is detected in both the CA and CC (Fig. 3F vs. 3G), but BaboA-GFP signals are very weak (Fig 3H). The dramatic reduction of this isoform from larval to adult stages predicts that BaboA might be also involved in a stage-specific aspect of tissue function and/or metamorphosis-related changes. To our surprise, we detected BaboB-GFP expression in a few CA cells and some esophagus and proventriculus cells in about 28% of specimens (Fig 3I). BaboC-GFP signals are detected weakly in the adult CA but strongly in the adult CC, again reminiscent of the larval pattern (Fig 3J). In summary, we found BaboA and BaboC isoforms in all the RG-associated endocrine systems from juvenile to adult stages. While BaboC expression is similar between the two stages, BaboA levels are dramatically reduced at maturity. This signifies a sophisticated splicing strategy in use to regulate isoform expression patterns and levels of the Babo isoforms in cell-and stage-specific manners.

As expected from the rescue results (Fig. 2), our results demonstrated that the C-terminally tagged Babo proteins are properly targeted to the plasma membrane of the RG cells (Fig 3B). In contrast, the GFP-conjugated isoforms spread in the cytoplasm, meaning that the tag disrupts membrane targeting of the Babo isoforms. We expect that such mis-targeting disables their functioning as receptors. However, the mis-targeting does not seem to be universal. For instance, BaboB-GFP proteins are found on the plasma membrane in adult gut cells, suggesting that membrane targeting of this reporter isoform might be less disrupted or normal in these cells (Fig 3I). These observations are consistent with our rescue assays.

### Detection of GFP-tagged Babo isoforms in the larval body wall tissues

Larval body wall harbors the muscle, the epidermis, and the oenocytes. Prior work has revealed that Actβ signaling from the motor neurons to muscles is a key to muscle development during the 3rd instar stage [12, 33]. Hence it could be that BaboB is responsible for the Actβ signal; however, BaboB expression has not been resolved in the muscles as well as other the body wall tissues. To address this, we stained the body walls of the GFP-tagged *babo* isoform lines. The GFP immunostaining gives none or a very low signals in the muscle and epidermis of *w^1118^* (Fig 4Aa-a¢). In the muscle, BaboB-GFP is the highest among the three isoforms and the protein is enriched at the postsynaptic area delineated by anti-Discs large (Dlg) staining (Fig 4Ac-c¢). BaboA-GFP and BaboC-GFP are also expressed but weakly in the muscle and enriched at the synaptic area as well (Fig 4Ab-b¢, 4Ad-d¢). Of note, synaptic enrichment does not necessarily mean a specific targeting of the proteins to the synapse. The postsynaptic area of *Drosophila* larval muscle bears dense membrane folds called subsynaptic reticulum (SSR) that oftentimes nonspecifically traps proteins in its complex structure. For example, cytosolic GFP expressed in the muscle is also enriched at the synaptic area delineating the peculiar neuromuscular junction (NMJ) shape (data not shown). To verify the highest expression of BaboB, we expressed *baboB-miRNA* in the muscle together with the *pan-babo-GFP* construct serving as a reporter. Pan-Babo-GFP is detected in the muscle, as expected, of control and is enriched at the synaptic area (Fig 4Ae-e¢), but muscle-specific *baboB-*KD (*mef2*>*baboB-miRNA*) significantly decreases the pan-Babo-GFP abundance (Fig 4Af-f¢), verifying that BaboB is the major isoform expressed in the muscle, consistent with the observation that Actβ-BaboB signaling controls muscle size [12]. The observations together with rescue assays also led us to speculate that the GFP tagged isoforms are functional receptors at least in the muscle because of their NMJ localization.

**Fig 4.**
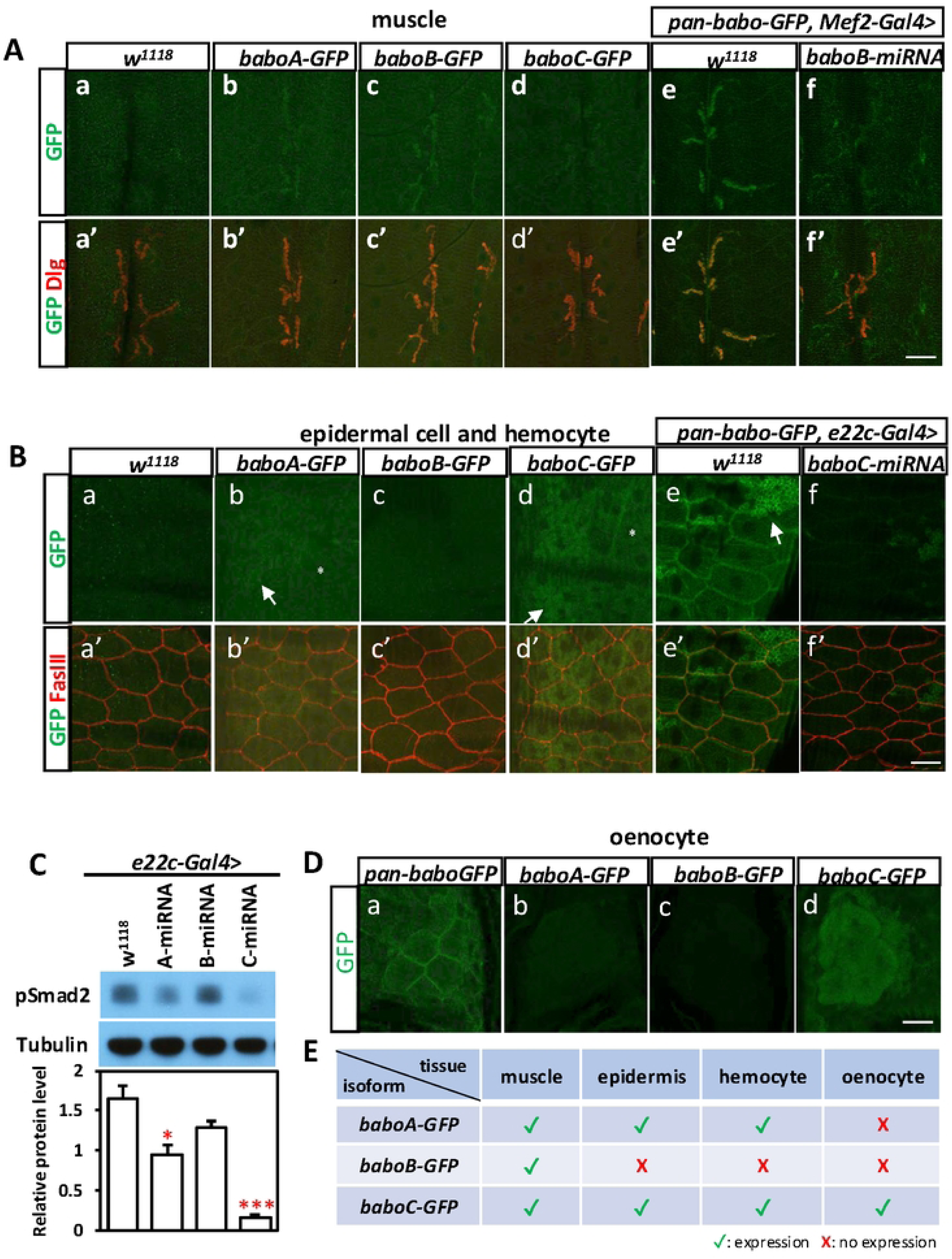
Expression of GFP-tagged Babo isoforms in the larval body wall. (A) Anti-GFP and anti-Dlg immunostainings on the muscles of *w^1118^* (a-á), *baboA-GFP* (b-b’), *baboB-GFP* (c-ć) and *baboC-GFP* (d-d’). The same staining was done to label pan-Babo*-*GFP at the synapse of control (e-é) and *baboB*-KD (*Mef2*>*baboB-miRNA*) (f-f’). (B) Anti-GFP and anti-FasIII staining on the epidermis of *w^1118^* (a-á), *baboA-GFP* (b-b’), *baboB-GFP* (c-ć), and *baboC-GFP* (d-d’). The same staining was used to detect pan-Babo-GFP on the control (e-é) and *baboC-*KD (f-f’). Hemocytes are indicated by arrows and nuclei by asterisks. (C) Immunoblot images of pSmad2 and Tubulin and a quantification graph showing relative protein value of each isoform. Samples were prepared from body walls expressing miRNAs of *babo* isoforms using *e22c-Gal4*. *P<0.05 and ***P<0.001 from one-way ANOVA followed by Dunnett’s multiple comparisons test in which each genotype was compared to *w^1118^.* (D) Anti-GFP staining on the oenocytes of indicated genotypes. (E) A summary of isoform expression in body wall tissues. Scale bars, 40 μm.

To investigate epidermal expression of the GFP-tagged isoforms, larval body wall fillets were co-stained with anti-GFP and anti-FasIII. The latter labels the boundary of epidermal cells. Unlike in muscle where BaboB-GFP exhibits highest expression, the epidermal cells do not show detectable BaboB-GFP (Fig 4Bc-c¢). Instead, this tissue contains BaboA-GFP and BaboC-GFP with higher expression given by BaboC-GFP (Fig 4Bb and 4Bd). Both are detected in the cytoplasm outside the perinuclear region (asterisks in Fig 4Bb and 4Bd), whereas pan-Babo-GFP clearly delineates the cell boundary (Fig 4Be-e¢). The highest expression of BaboC-GFP is further demonstrated by substantial depletion of pan-Babo-GFP by *baboC-miRNA* expressed in the epidermal cells via *e22c-Gal4* (Fig 4Bf-f¢). Next, we performed Western blot analysis to assess the functional consequence of each *babo* isoform KD in the epidermal cells. As presented in Fig. 4C, *e22c*>*baboC-miRNA* leads to the most significant decrease in the pSmad2 level, which is a readout of canonical Activin signaling. Expression of *baboA-miRNA* also decreases the pSmad2 level but to a lesser extent, while *baboB-miRNA* is ineffective. These results are consistent with the differential expression levels of Babo isoforms in the epidermal tissue. In addition to epidermal cells, BaboA-GFP and BaboC-GFP are observed in hemocytes that are occasionally fixed with body wall tissues (arrows in Fig 4Bb, 4Bd, 4Be).

Oenocytes consisting of large clusters of secretory cells positioned along the lateral body wall are known to play an important role in lipid metabolism [34, 35]. Pan-Babo-GFP is detected in the oenocytes with an enrichment at the plasma membrane (Fig 4Da). BaboC-GFP is found in this cell type, while BaboA-GFP and BaboB-GFP expression is undetectable (Fig 4Db-d). As in the epidermal cells, oenocytic BaboC-GFP does not delineate the cell boundary, indicating its cytoplasmic targeting (Fig 4Dd). The expression of GFP-tagged Babo isoforms in larval body wall is summarized in Fig 4E.

### Two Babo isoforms expressed in the CNS

Loss-of-function *babo* mutants exhibit a small brain phenotype, largely due to defects in promoting proliferation of neural stem cells and NE-to-NB conversion, suggesting important roles for this receptor in regulating these cellular events during postembryonic neurogenesis [10, 36]. To gain further insight into Babo’s neural functions, we examined pan-Babo-GFP expression in the CNS at WL3 stage. Babo expression is manifested broadly in the brain and the ventral nerve cord (VNC) (Fig 5A-5D). To resolve if this gene is expressed in two neural stem cells, NBs, and NEs, pan-Babo positive cell types were double-labeled with anti-Dlg, which marks both cell types as well as ganglionic mother cells (GMCs) and precursor cells in the postembryonic CNS (Fig 5Ai-5Di). Notably, all Dlg-labeled cells also contain pan-Babo-GFP (Fig 5Aii-5Dii). The patterns are consistent with ones of *babo^fTRG00444.sfGFP-TVPTBF^* flies, which carry multi-tags including GFP at the C-terminal like pan-Babo-GFP (S1 Fig).

**Fig 5.**
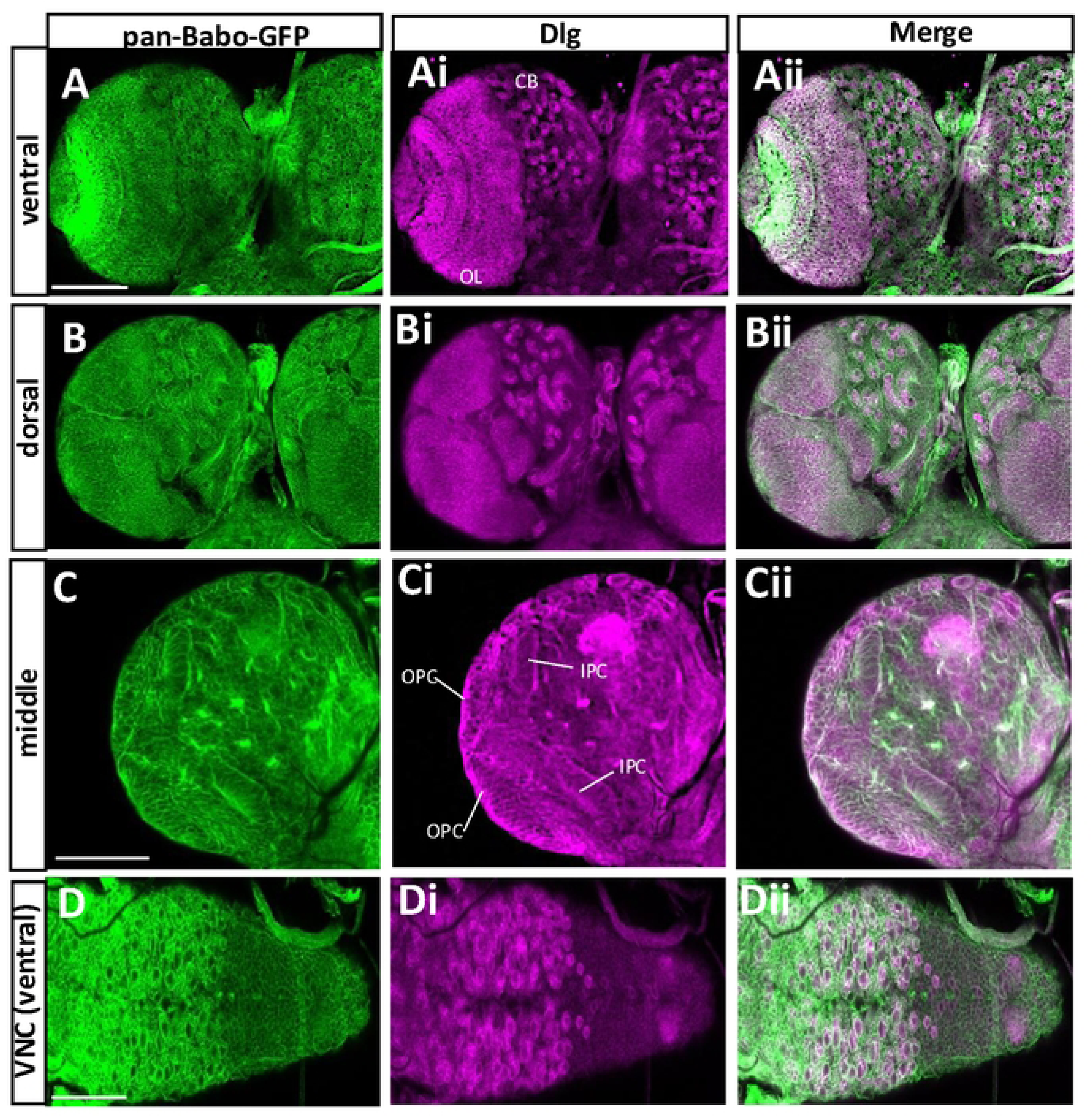
Expression patterns of pan-Babo-GFP in the CNS at WL3. The CNSs of pan-*babo-GFP* (n=14) were processed with anti-GFP (green) and anti-Dlg (magenta). Ventral (A-Aii), dorsal (B-Bii), and middle sections (C-Cii) of a brain lobe are shown. (D-Dii) Ventral side of a VNC. CB, central brain; OL, optic lobe; OPC, outer proliferation center; IPC, inner proliferation center; Scale bars in A and D, 100 μm, and in C, 50 μm.

Next, we wanted to determine which isoforms are accountable for the pan-Babo pattern and found that BaboA is the most prevalent and strongly expressed one (Fig 6A-Ai and 6B-Bi) and moderate levels of BaboC that are significantly lower than BaboA (Fig 6B-Bi vs. 6C-Ci) and no detectable BaboB (Fig 6D vs. 6E). These results imply that pan-Babo represents a sum expression of BaboA and BaboC. These two isoforms are also detected in the adult CNS (S2 Fig); however, their expression levels are not as dramatically different as in the larval stage.

**Fig 6.**
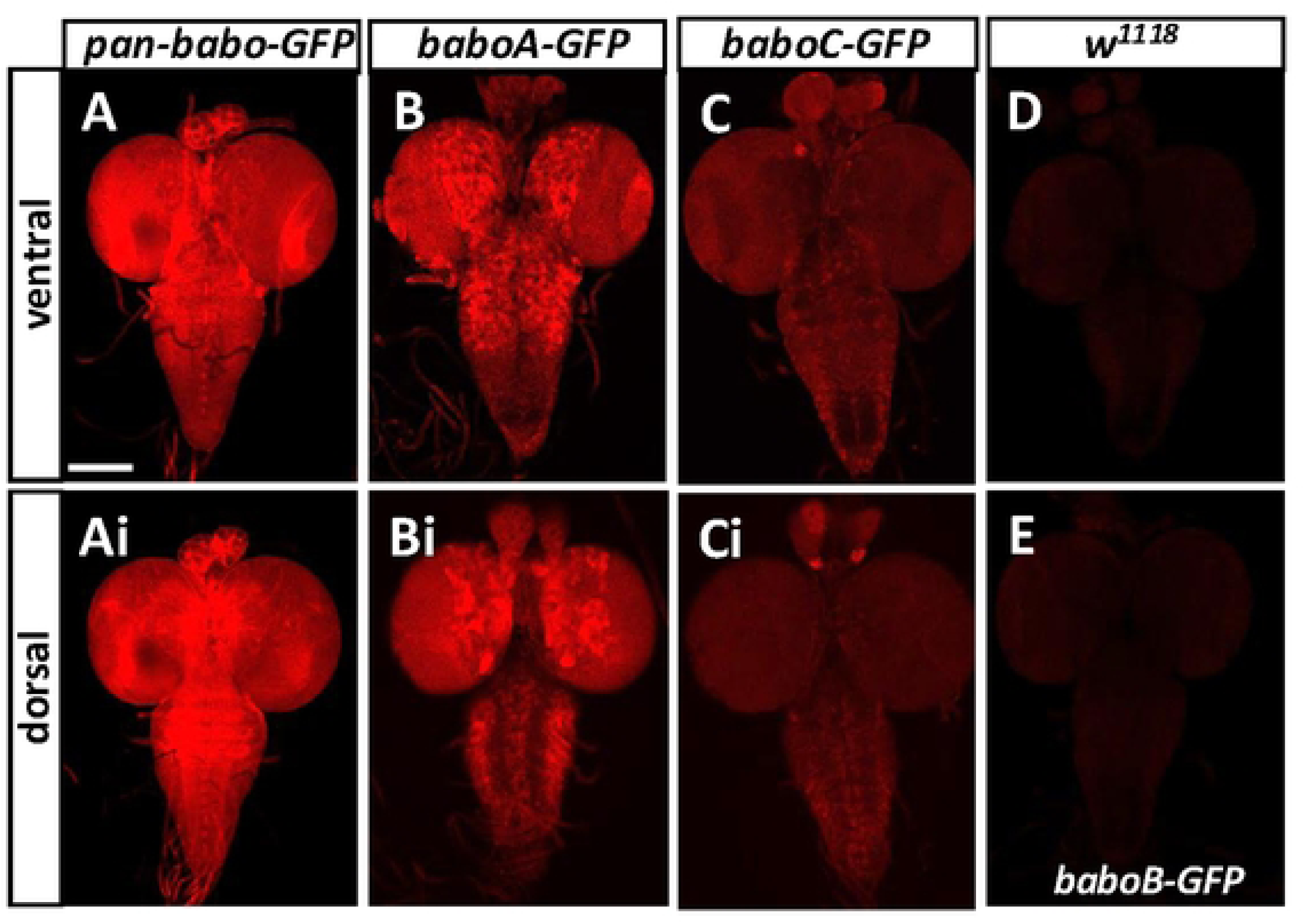
Detection of Babo isoforms in the larval CNS at WL3 stage. The larvae are homozygous for each transgene. All genotype samples were processed simultaneously and imaged under the same condition. (A-E) Ventral sides of representative CNSs are shown for expression of (A) *pan-babo-GFP* (n=14), (B) *baboA-GFP* (n=24), (C) *baboC-GFP* (n=24), (D) *w^1118^* (n=16), (E) *baboB-GFP* (n=16). (Ai-Ci) Dorsal side of the same CNSs as in A-C. Scale bar: 100 μm.

### BaboC expression in the CNS

Closer examination of BaboC-GFP expression revealed distinctive patterns in the larval CNS (Fig 7A-7K, 7Fi-7Ji). This isoform is found in several neuronal clusters of one to eight cells in the ventral protocerebrum and deutocerebrum (Fig 7C and 7D). Based on their number and position, one of these clusters appears to be the well-characterized insulin-producing cells (IPCs) that are shown to produce three different Drosophila insulin-like proteins (Dilp2, 3, and 5) [37]; indeed, these cells are co-labeled with anti-Dilp2 (Fig 7L-7N). BaboC-GFP expression is additionally found in numerous neurons in the suboesophageal, thoracic, and abdominal ganglia where a large population of motoneurons are present, including ones in the dorso-median region (Fig 7C, 7G, 7H, 7K). Glial cells also express this isoform, as confirmed by co-staining with a glia marker, anti-Repo (Fig 7Ai-7Ji). As in the RG cells, the BaboC-GFP is cytoplasmic in neuronal cells. However, to our surprise, this is not the case for glial cells. The tagged proteins are mainly found in glial processes, indicating their membrane targeting in glia. It is not clear why neuronal and glial cells handle the trafficking of BaboC-GFP protein differently. Regardless of its nature, the presence of BaboC-GFP in the glial processes permits us to resolve at least two glial subtypes, perineurial glia (pg) and cortex glia (cg), which are identifiable by their process patterns and location [38–41]. Judging from the patterns, BaboC is expressed in the pg that is one of two surface glial subtypes which together form the blood brain barrier (BBB) and in the cg that is found in the cortex beneath the surface and displays honeycomb-like processes wrapping around individual NB lineages (Fig 7A-7E, 7Ai-7Ei). There are two neuropil glia subtypes located deep inside the CNS that are closely associated with neuronal axons, and this isoform is present in the neuropil glial cells located along with the longitudinal lateral track of the VNC (Fig 7Ii-7Ji).

**Fig 7.**
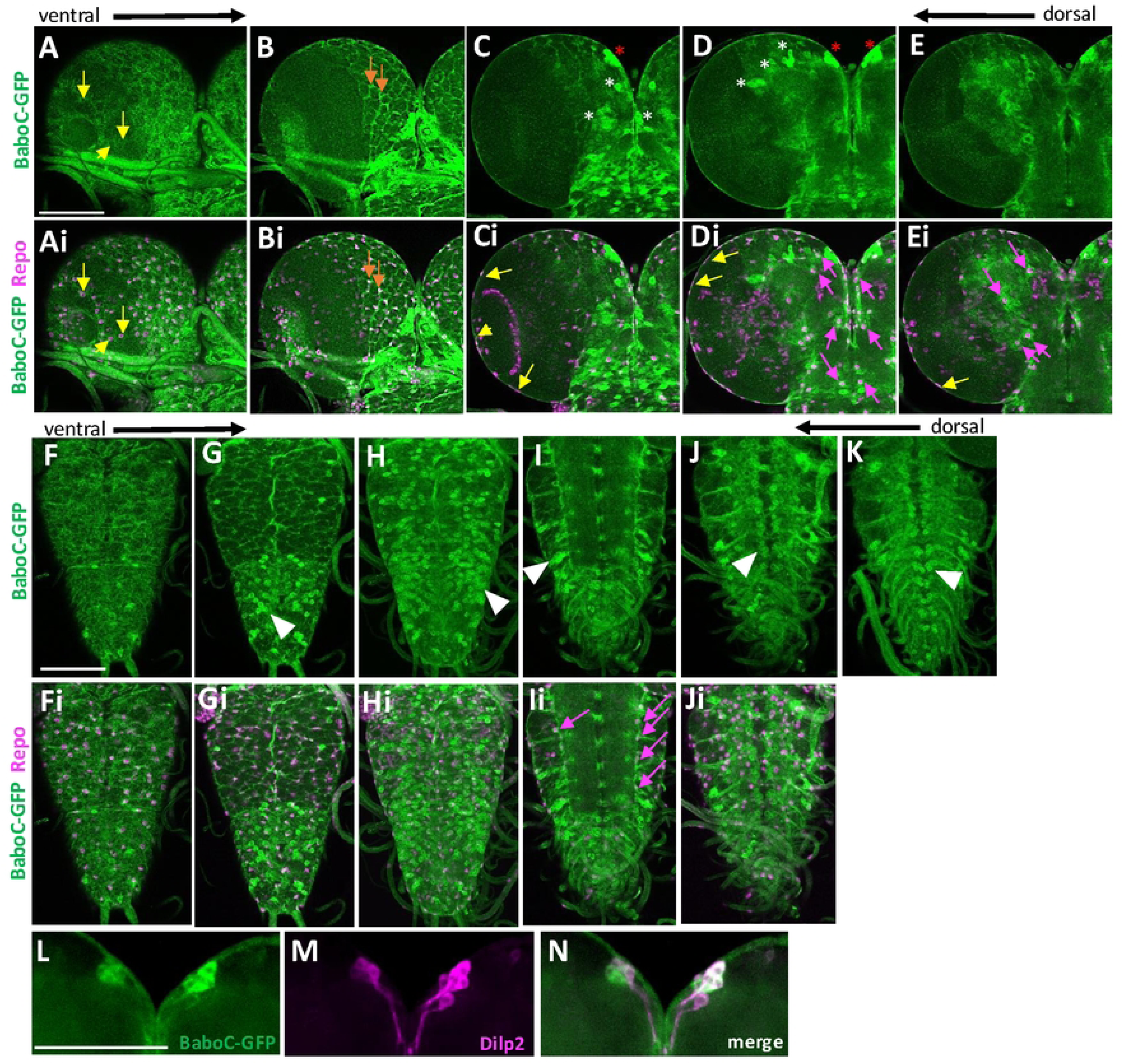
BaboC-GFP expression in larval neurons and glial cells. CNSs (n=14) of *baboC-GFP* larvae at WL3 stage were processed for double-labeling with anti-Repo (magenta) and anti-GFP (green). From ventral to dorsal optical sections of a brain lobe showing GFP alone (A-E) and with Repo (Ai-Ei). Different glial cell subtypes are indicated by arrows: yellow (surface glia), orange (cortex glia), and magenta (astrocyte-like or ensheathing glia). Asterisks indicate Repo-negative neuronal clusters. From ventral to dorsal sections of a VNC showing GFP alone (F-J) and with Repo (Fi-Ji). Expression of BaboC-GFP in a large population of Repo-negative neurons and dorsal-median motoneurons is indicated by arrowheads. (K) A composite of three dorsal z-sections to show dorsal-median motoneuron clusters. (L-M) BaboC-GFP (L) in the Dilp2-producing median neurosecretory cells (M,N). Scale bars in A, F, and L, 50 μm.

We also asked if this isoform is expressed in NBs and NEs by double-labeling with anti-Dlg. No apparent presence of this isoform is found in the NBs of the central brain (CB) and their lineage cells nor in NEs and medulla NBs of the OL (Fig 8). Instead, we found GFP-marked glial processes enveloping around NB lineages as well as the proliferation centers and medulla NB lineages in the OL. They are presumably the cg subtype, which is reported to wrap around the outer proliferation center (OPC) and medulla NB lineages in a manner similar to the CB ones [42].

**Fig 8.**
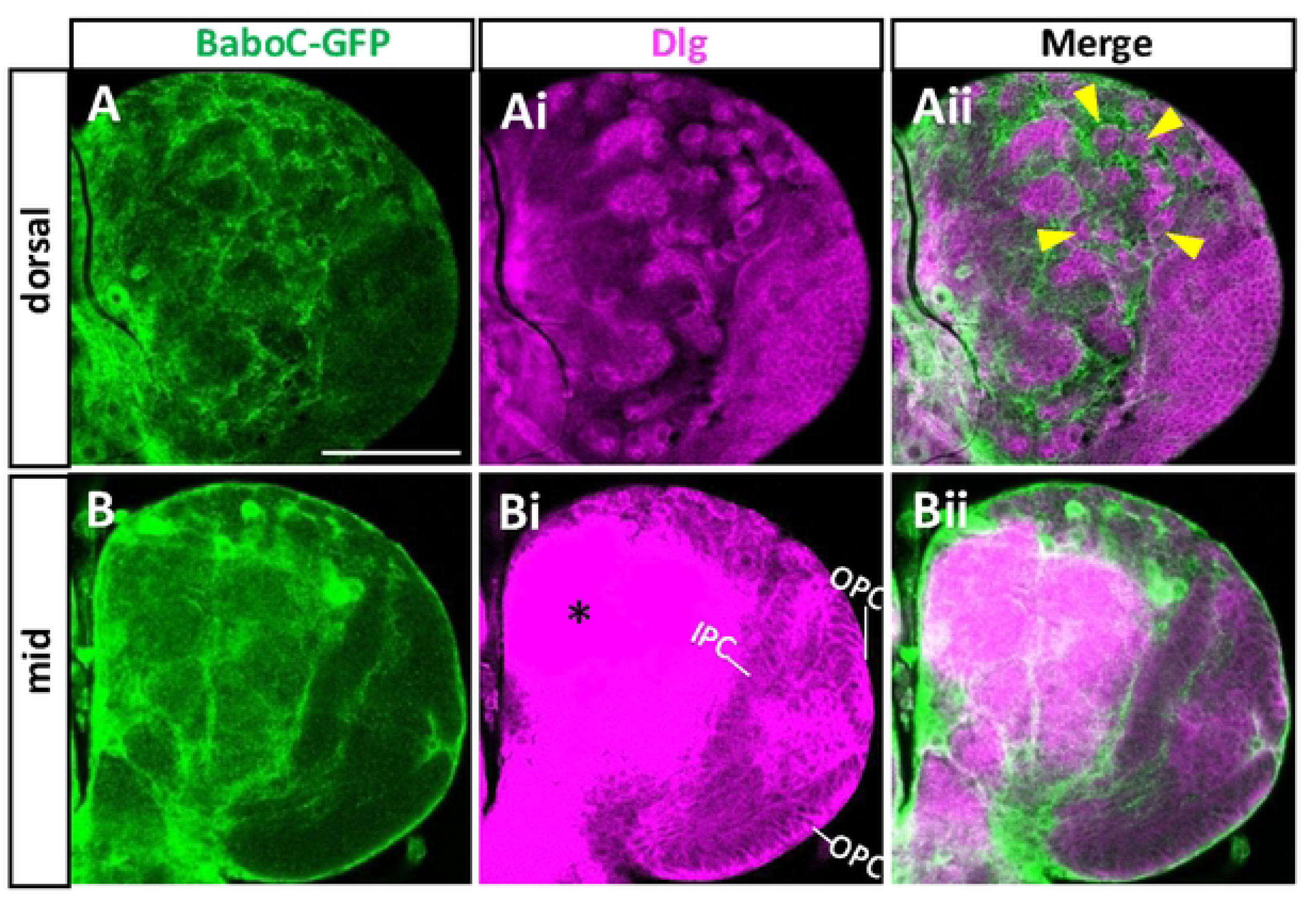
BaboC-GFP is absent in the NBs and NEs. Anti-Dlg (magenta) labels NBs and NEs of larval CNS. (A-Aii) NBs in the dorsal central brain lack BaboC-GFP expression as indicated by arrowheads (n=14). (B-Bii) BaboC-GFP expression is absent in NEs of both IPC and OPC. An intense Dlg staining (*) in the neuropil. A scale bar in A, 50 μm.

Previous study has shown that the TGF-β signaling mediated by Myo/BaboA/Smad2 operates autonomously to upregulate EcR-B1 expression levels in the MB neurons [7]. We confirmed the presence of both pan-Babo-GFP and BaboA-GFP in them (Fig 9A-Aii, 9B-Bii). Surprisingly, BaboC-GFP is also detected in these neurons (Fig 9C-Cii), indicating that the pan-Babo-GFP expression in MB neurons represents both isoforms (Fig 9A-Aii).

**Fig 9.**
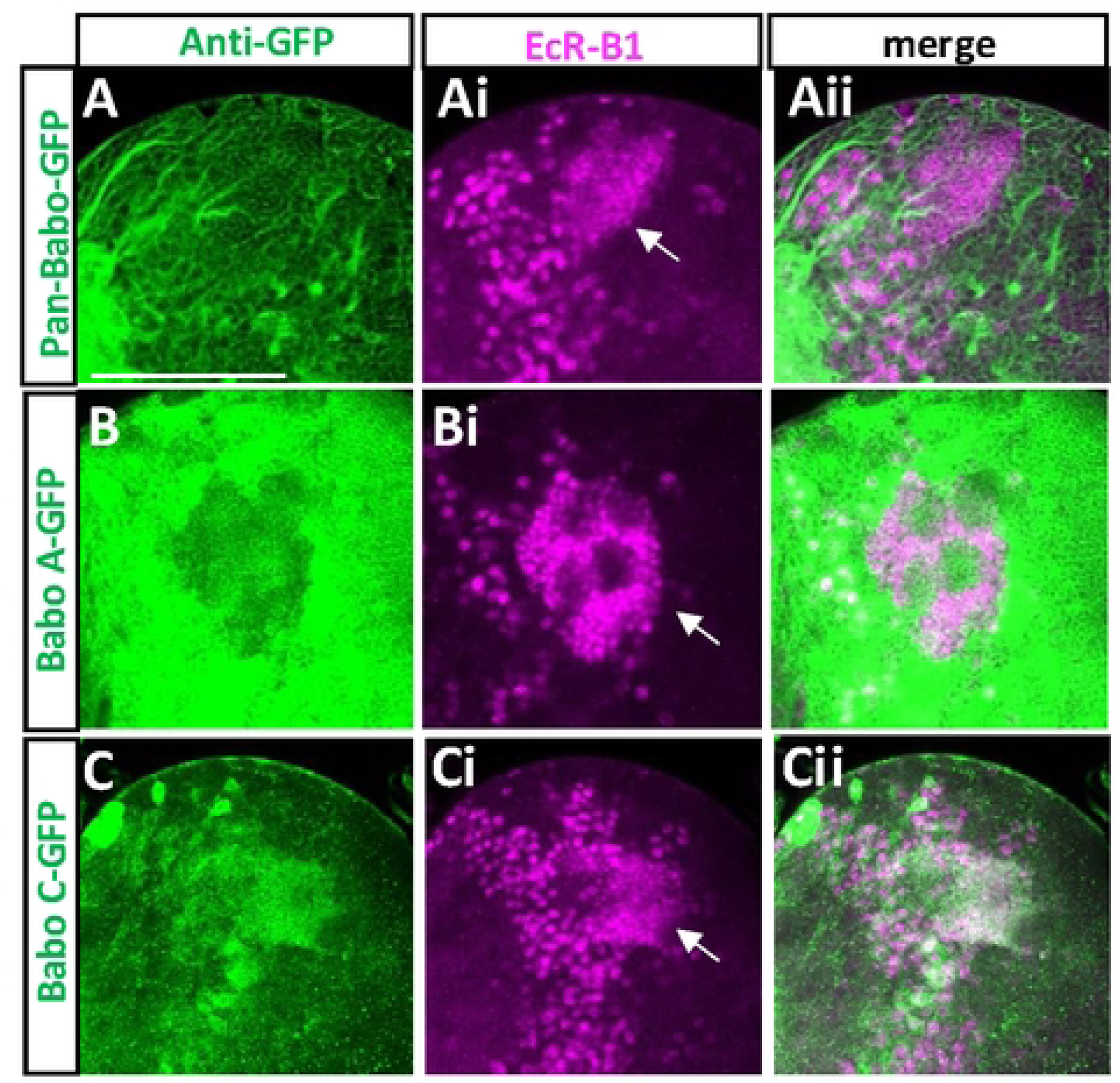
Expression of BaboA- and BaboC-GFP in the mushroom body (MB) neurons. Anti-EcR-B1 (magenta) was used to label MB neurons indicated by arrows. (A-Aii) pan-Babo-GFP (n=7). (B-Bii) BaboA-GFP (n=14). (C-Cii) BaboC-GFP (n=14). A scale bar in A, 50 μm.

### Neural Cell types expressing BaboA

Our previous genetic data showed that BaboA is the major isoform functioning for postembryonic neurogenesis [10]. Consistently, BaboA-GFP is found in a wide range of cell types of the larval CNS (Fig 10). It is located exclusively in the cytoplasm of these neural cells, indicating the GFP insertion disrupts the membrane-targeting of the tagged isoform in them. Another observation is its variable levels depending on cell type. The most intense staining is seen in many of the densely clustered NB lineage cells in the CB and thoracic ganglia (Fig 10A-10C). Clusters of larval motoneurons, particularly in the dorso-median region of the thoracic and abdominal ganglia also show strong signals (Fig 10D). Most cell types in the OL and abdominal ganglia show from intermediate to low levels of BaboA (Fig 10A-10C). This isoform is also detected in most, if not all, of larval neurons that are generated during embryonic neurogenesis (S3C Fig).

**Fig 10.**
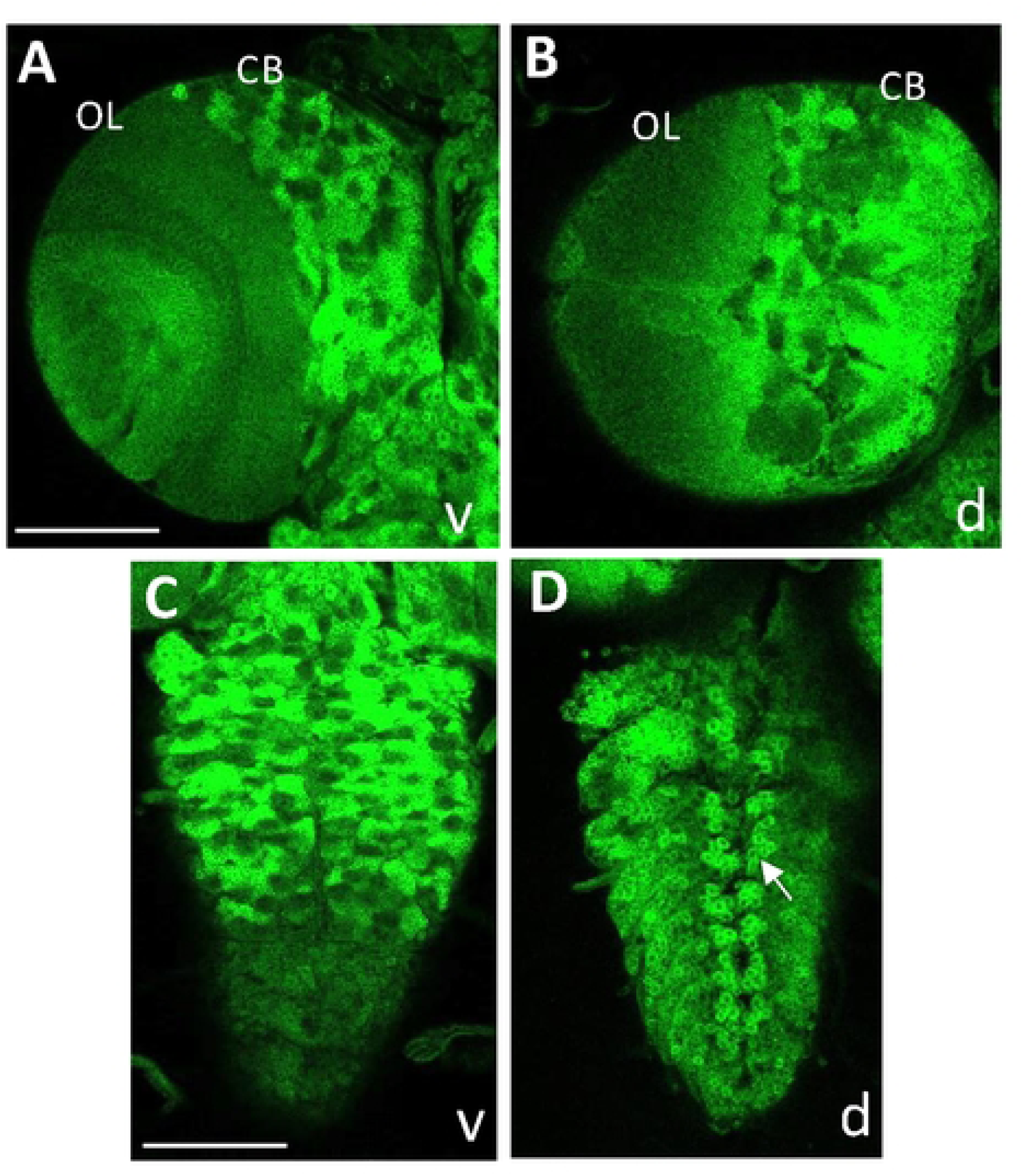
Expression of BaboA-GFP in the larval CNS at WL3 stage. The larvae are homozygous for *baboA-GFP* (n=14). BaboA-GFP expression is shown in ventral (v) and dorsal (d) sides of a brain lobe (A,B) and a VNC (C,D). A cluster of dorso-median motoneurons is indicated by an arrow in (D). Scale bars in A and C, 50 μm.

To further ascertain BaboA expression in NEs and NBs, *baboA-GFP* brains were double-labeled with ant-Dlg or anti-Miranda (Mira). While Dlg labels all cell types of postembryonic neurogenesis, Mira labeling is specific to NBs and recently born GMCs. Both proteins are detected on the plasma membrane. We found BaboA-GFP expression in all cell types positive for both Dlg and Mira (Fig 11). Interestingly, individual precursor cells in the CB and thoracic ganglia display either high or low levels of BaboA expression, which leads to a distinct pattern for each NB lineage (Fig 10A-10C, 11Ai-11Bi, S3Ai-Ei). Meanwhile, precursor cells of the OL express BaboA rather weakly but more-or-less evenly. These observations suggest that this isoform might play diverse roles during postembryonic neurogenesis.

**Fig 11.**
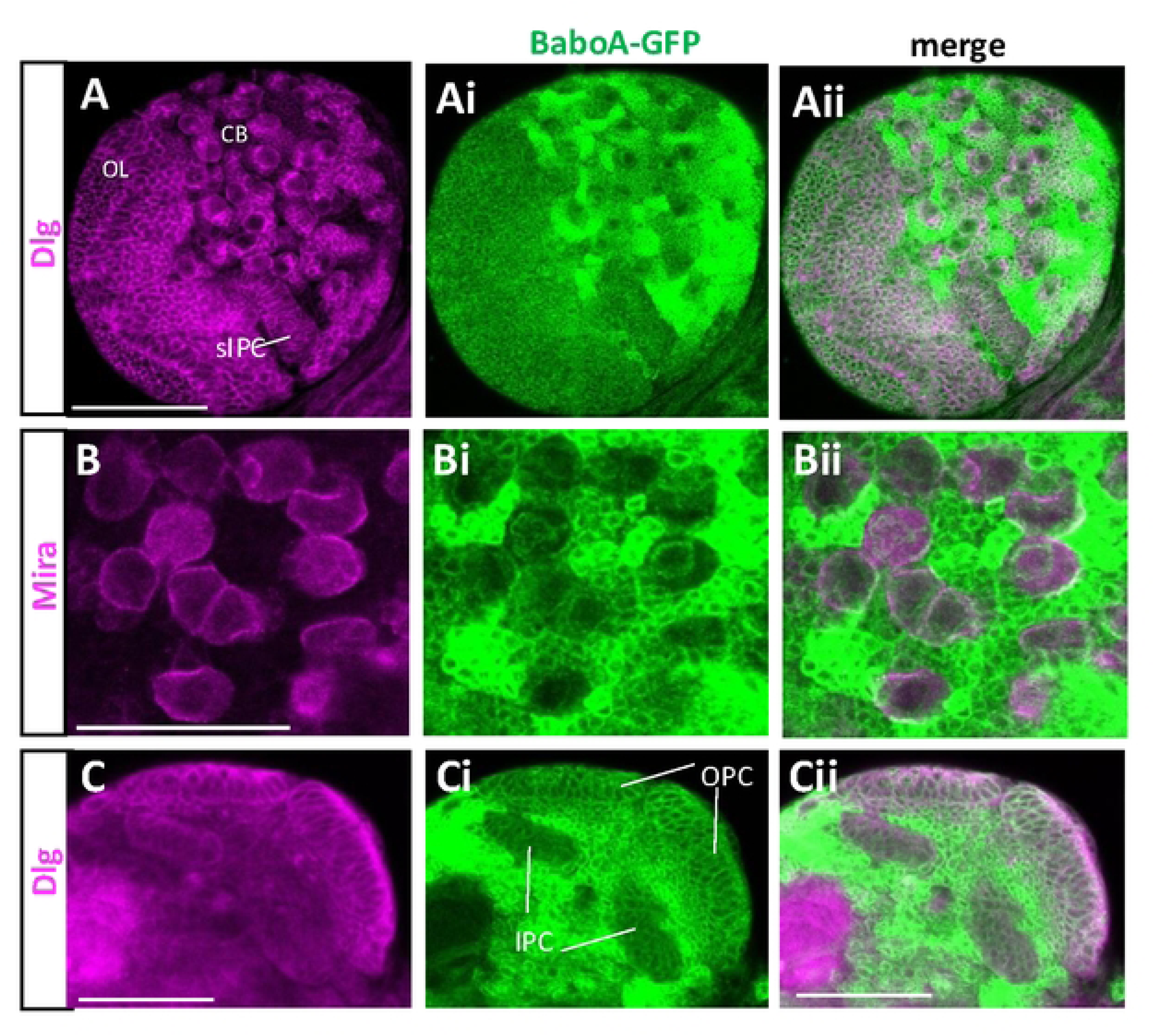
BaboA-GFP expression in the NBs in the CB and NEs and medulla NBs in the OL of the L3 brains. (A-Aii) BaboA-GFP and Dlg is shown in dorsal side of a brain lobe. Surface IPC (sIPC) is indicated in A. Dlg and Mira proteins (magenta) are detected on the cell membrane and BaboA-GFP, in cytosol. (B-Bii) Detection of BaboA in all Mira-labeled NBs in the CB. (C-Cii) Detection of Dlg and BaboA-GFP in the NEs of both OPC and IPC, medulla NBs, and medulla precursor neurons at 96 h AEL. Scale bars in 50 μm.

A precursor cell in the CNS undergoes an initial decision process that determines its development to either neuronal or glial path. To see if BaboA expression levels are somehow linked with this major decision-making step, we double-labeled the precursor cells with a pan-neuronal marker Elav that marks the neuronal status. A large population of precursor cells in individual NB-lineages are Elav-positive, suggesting that the neuronal fate determination event takes place soon after their birth or by WL3 stage (Fig 12A-Aii). Since BaboA-GFP expression is detected in all precursor cells either weakly or strongly, we focused on those with high levels of BaboA-GFP expression. These cells are generally positive for Elav. However, we also found intense BaboA-GFP signals in small populations of Elav-negative cells suggesting that these are either prior to neuronal fate determination or glia-fated cells. BaboA-GFP is also expressed in loosely distributed numerous Elav-positive neurons in the median regions of the protocerebrum and subesophageal ganglia which fit the characteristics of differentiated larval neurons based on their soma size and cytoplasmic volume, while precursor cells are tightly clustered small cells within a cytoplasmic strip (Fig 12A-Aii).

**Fig 12.**
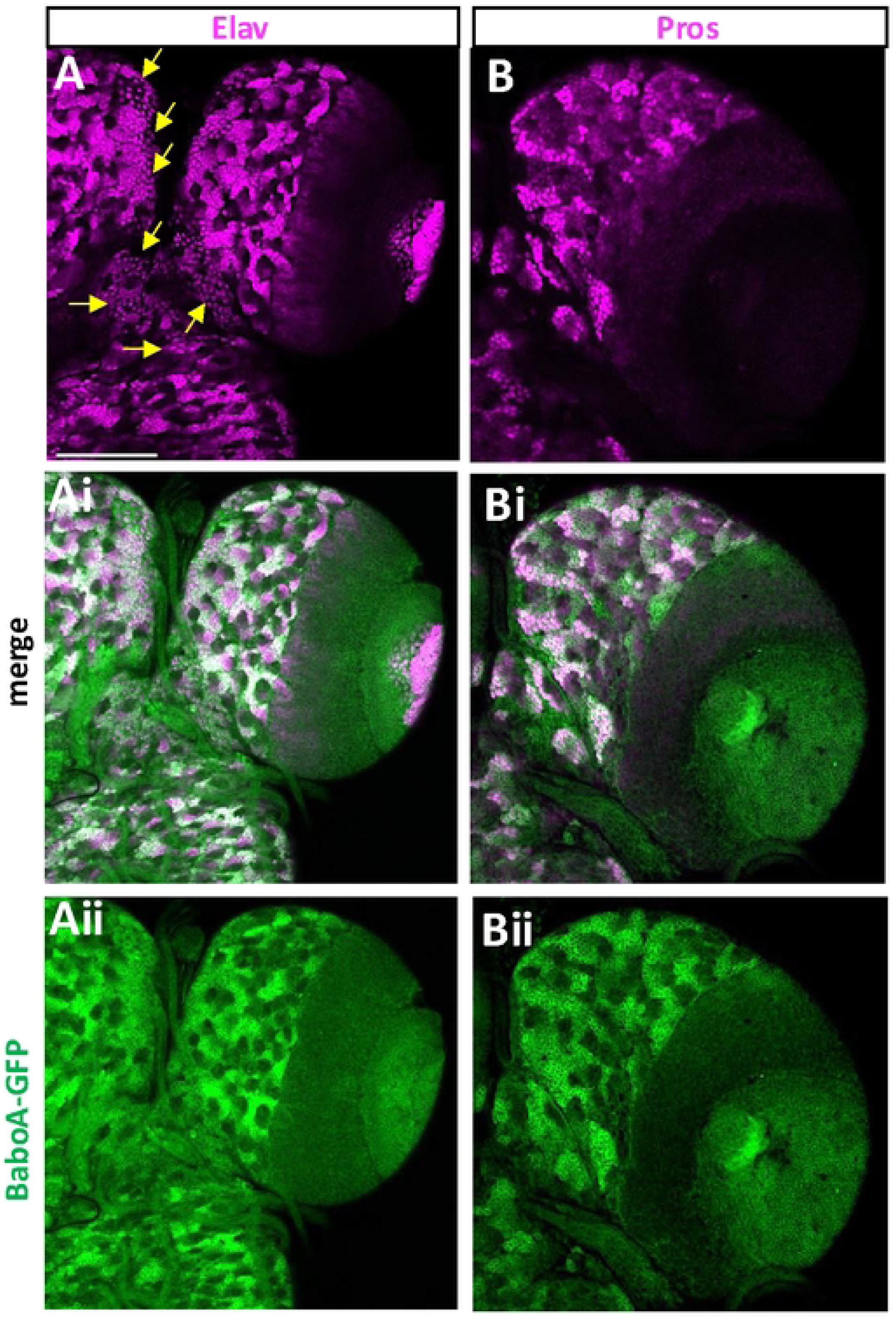
BaboA-GFP expression in the cell types positive for Elav and Pros in the brain at WL3 stage. (A-Aii) BaboA-GFP in Elav-positive mature and immature neurons (n=14). Arrows indicate loosely distributed mature larval neurons. (B-Bii) BaboA-GFP in precursor cells showing both strong and weak expression of Pros in individual NB lineages in the CB (n=8). A scale bar in (A), 50 μm.

Prospero (pros) is present in all postmitotic cells of NB lineages in the larval CB, but its expression levels are low before cell-type specification, which event follows neuronal determination, and become higher after this state [43, 44]. Notably, there are more Elav-positive neurons than ones with high levels of Pros at WL3 stage (Fig 12A and 12B). Because of intense Pros expression marking differentiated neurons in each NB lineage, we reason that many Elav-positive neurons with weak Pros signals have not fully undergone the cell-type specification process by this stage. Most postmitotic cells with high levels of Pros also express BaboA-GFP strongly (Fig 12B-Bii). However, the opposite is not always the case. These results indicate that high levels of BaboA expression are seen in both fated and non-fated cells.

### BaboA expression in glial cells

Given the wide distribution of BaboA-expressing cell types, we questioned if glial cells also express this isoform. Despite clear Repo labeling of glial nuclei, it was a challenge to determine if these cells are also positive for BaboA-GFP (Fig 13A-Aii). Thus, we focused on glial cells located on the OL surface and on the dorsal side of the VNC. On the OL surface, two kinds of glial cells are positive for BaboA expression based on nuclear size (Fig 13B-Bii). The glial cells with large nuclei seem to fit characteristics of the surface glia [39]. We also found BaboA-GFP-positive glial cells in the dorso-lateral region of the VNC (Fig 13C-Cii). To see if they are astrocyte-like glia (ag), we co-labeled with anti-Pros, which nuclear expression is found in the ag located in the VNC [45]. Pros-positive ag cells indeed contain cytoplasmic BaboA-GFP (Fig 13J-13L). In summary, BaboA isoform is expressed at least in two glial subtypes.

**Fig 13.**
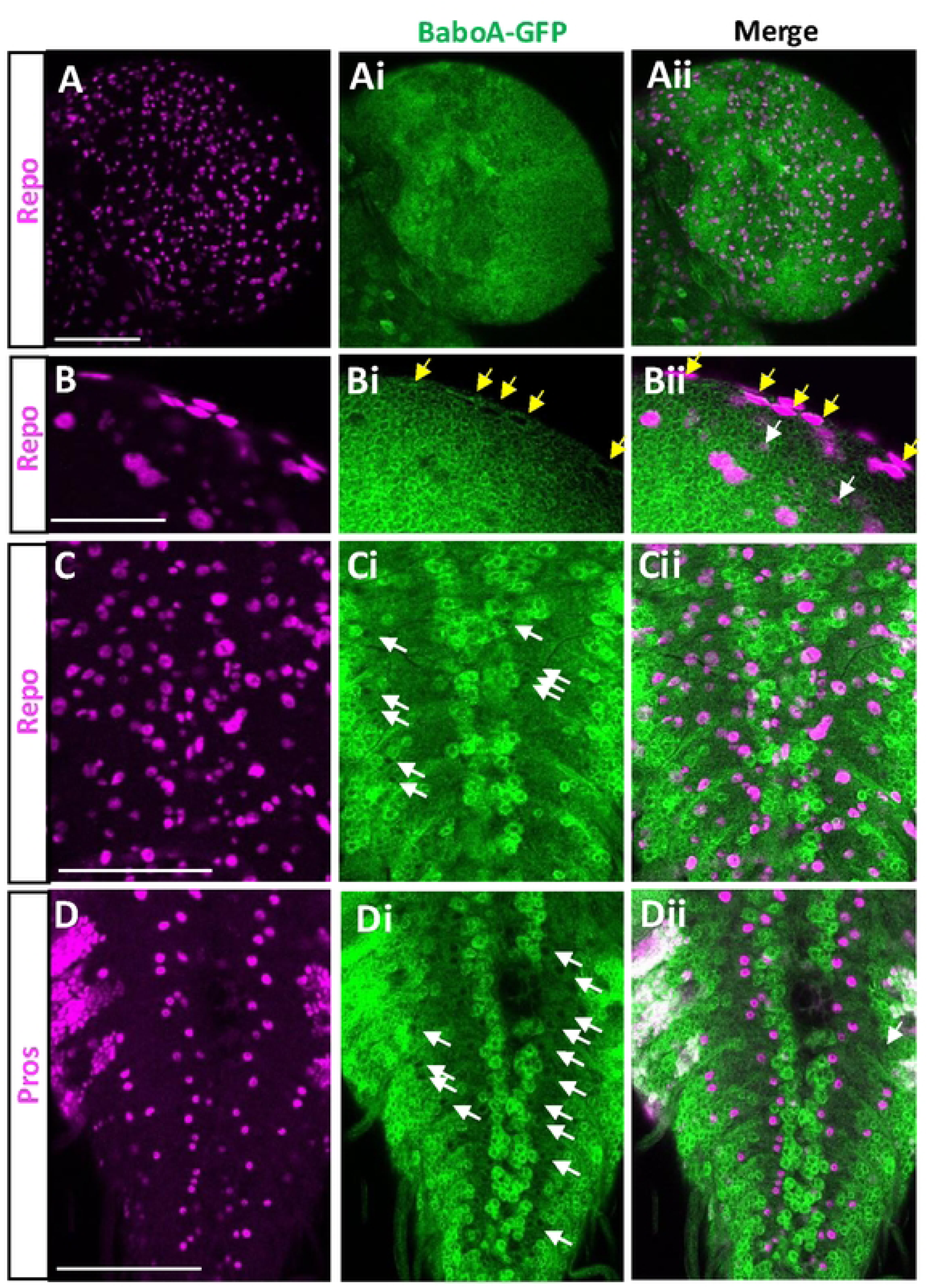
Glial expression of BaboA-GFP in the brain at WL3 stage. (A-Aii) A dorsal side of a brain lobe showing broad distribution of Repo-positive glial cells (magenta) and BaboA-GFP (green). (B-Bii) BaboA-GFP in surface glial cells (yellow arrows) and small-sized glial cells of the OL (white arrows). (C-Cii) BaboA-GFP in a few Repo-positive glial cells (arrows) in the dorsal side of a VNC. (D-Dii) BaboA-GFP in astrocyte-like glial cells labeled by Pros (magenta). Scale bars in A, C, and D, 50 μm, in B, 25 μm.

### BaboA functions for peptidergic neurons

Although most *baboA-indel* mutants die during prepupal and early pupal development, a small percentage of them manage to develop to the pharate stage [10]. This permits us to examine if BaboA function is also required for generation of adult-specific neurons, cell fate determination, and/or their differentiation. Neuropeptides are convenient markers for differentiated neurons and each neuropeptide is produced by distinctive neuronal groups. We chose four neuropeptides, neuropeptide F (NPF), pigment dispersing peptide (PDF), corazonin (CRZ), and drosulfakinin (DSK), for the following reasons: availability of specific antibodies, and well-characterized neuroanatomical features of both persisting larval and adult-specific neurons [46–49]. We explored if their expression patterns and/or neuroanatomical features of the neurons are affected in the pharate brain of *baboA-indel* mutants. All larval clusters of the peptidergic neurons are present in *baboA-indel* pharates, but their overall axonal features closely resemble larval ones (Fig 14). Since the remodeling of larval axonal projections to adult ones is a default event for the persisting larval neurons, the results indicate that the persisting peptidergic neurons fail to undergo the remodeling process in the absence of BaboA function. We also point out that the vCrz neurons, which are normally removed by PCD during the early phase of metamorphosis, remain visible in the mutant pharate (Fig 14R), consistent with this isoform’s proapoptotic role for these neurons [9]. Of interest, the undead vCrz neurons retain their larval projection patterns, which is different from what is seen when cell death is blocked by ectopic expression of a caspase inhibitor P35, or through elimination of cell-death genes. In these cases, the surviving vCrz neurons underwent axon remodeling process like other persisting larval neurons [50, 51]. Based on these observations, we propose that BaboA function is needed generally for the metamorphosis-associated remodeling processes of larval neurons including peptidergic ones.

**Fig 14.**
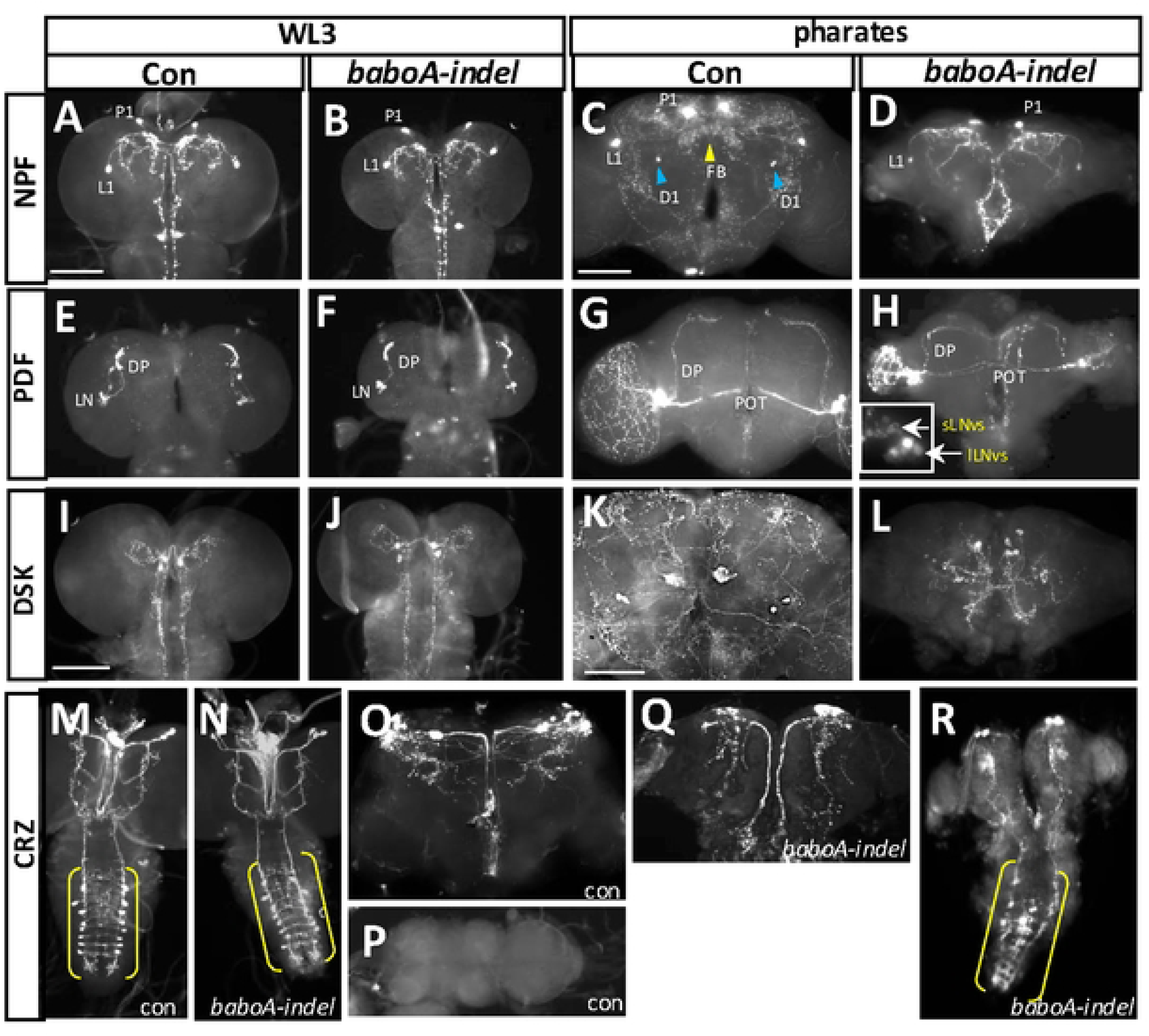
Various defects of peptidergic neurons in the absence of BaboA function. (A-D) NPFergic neurons in control (n=14) and *baboA* mutant (n=14) at WL3 stage (A,B), and in anterior sides of control (n=8) and *baboA* mutant (n=4) pharate brains (C,D). Blue arrowheads, a male-specific NPFergic neuronal group (D1) located near the antennal lobe; yellow arrowhead, axonal arborizations in the fan-shaped body (FB) derived from adult-specific neuronal group located in the posterior side that is common to both sexes. (E-H) PDFergic neurons (LN) and their dorsal projections (DP) in control (n=8) and *baboA* mutant (n=8) at WL3 stage (E,F), and those in pharate adult stage (G,H) of control (n=7) and *baboA* mutants (n=3). The posterior optic tracts (POT) from lLN_v_s are indicated. An insert in (H) is a magnified image to show sLN_v_s and lLN_v_s of the left lobe. (I-L) DSKergic neurons in the control (n=7) and *baboA* mutant (n=8) at WL3 stage (I,J), and those in pharates (K,L) of control (n=6) and *baboA* mutant (n=2). (M-R) CRZergic neurons in the control (n=7) and *baboA* mutant (n=7) at WL3 (M,N). vCrz neurons are in the brackets; in the brain and VNC of control pharates (n=7)(O,P). vCrz neurons are undetectable because of their PCD (P); in the brain and VNC of *baboA* mutant pharates (n=3) (Q,R). A few vCrz neurons (bracket) remain detectable in the *baboA* mutant. Scale bars, 50 μm.

There are five adult-specific NPFergic neuronal clusters; three male-specific and two sex-nonspecific ones [47]. None of them are found in *baboA*-indel pharate brains by anti-NPF (Fig 14C and 14D). However, this was not the case for the three other peptidergic neurons, since their adult-specific clusters are detected in the mutants (Fig 14G-14H; 14K-14L; 14O-14R). These results suggest a specific role played by BaboA for generation or cell fate determination of NPFergic adult-specific neurons but not for the other peptidergic ones. On the other hand, BaboA function appears to be involved in maintenance of high-level PDF expression in four adult-specific large latero-ventral neurons (lLN_v_s), which send axons through the posterior optic tract to innervate the opposite side of the optic lobe [46, 52]. PDF is expressed substantially high in all lLN_v_s in wild-type (Fig 14G), whereas it shows variable levels ranging from moderate to very weak in *baboA*-*indel* mutants (Fig 14H).

### BaboA function essential for development of the medulla neurons

The medulla region contains approximately 40,000 interneurons, representing the largest structure in the adult visual nervous system [53–57]. Since BaboA-GFP expression is detected in all medullar NB lineages, we were curious if BaboA plays any role in cell fate determination of the medulla precursor neurons. Scarecrow (Scro) is one of the temporal factors for medulla NBs, which is expressed in aged medulla NBs and continuously in their descendent neurons but not in OPC NEs and retinal photoreceptor neurons [18, 58, 59]. The adult medulla neuropil is organized in 10 layers (M1-M10) with repetitive columnar units oriented perpendicular to the medulla layers and display defined columns which match the individual ommatidia [60, 61]. Anti-24B10 (Chaoptin) stains retinal photoreceptor neurons (R1-R8) and their axons [62]. R1-R6 axons terminate together and form a lamina plexus, while R7 and R8 ones extend to the medulla field and their pattern displays the columnar organization but not layers at WL3 stage (Fig 15A). In the adult medulla, the chaoptin staining visualizes axon terminals of R7 and R8 at the M6 and M3 layers, respectively, and their axons are evenly spaced and their targeting sites mirroring individual columnar units (Fig 15C-Ciii). In contrast, the axon targeting patterns of R7 and R8 are disrupted in the larval optic lobe of *scro>baboA-miRNA* (Fig 15B). In adult, the patterns are even more severely compromised; the medulla lacks the chaoptin-labeled M3 and M6 layers and shows interruptions in even spacings of the running retinal axons (Fig 15D-Diii). Such defective chaoptin patterns show an uncanny resemblance to abnormal axonal tracts of the mCD8GFP-labeled mutant medulla neurons (Fig 15Eii-Eiii) [18]. These results suggest that *baboA*-depleted medulla neurons fail to generate typical neuropil structure, the M3 and M6 layers, and evenly spaced columnar organizations by the medulla neurons. Our prior study has shown that generation of the precursor neurons by medulla NBs are largely unaffected in *scro>baboA-miRNA* at the WL3 stage [10]. Taken together, we propose that BaboA function is required for either cell-type specification or differentiation of the medulla neurons, which is necessary for proper neuropil organization of the adult medulla system.

**Fig 15.**
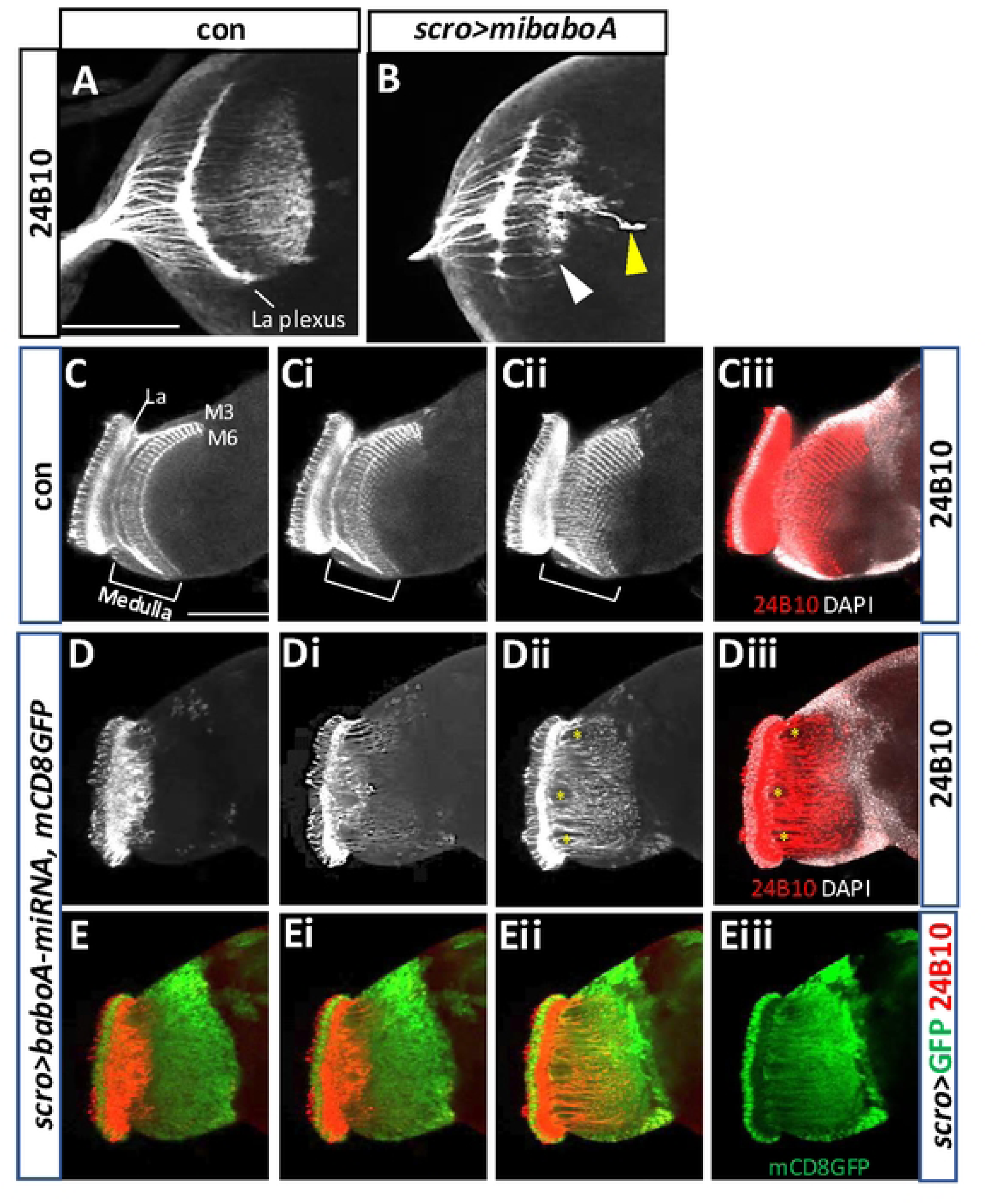
*baboA*-KD causes defective development of the medulla neurons. *scro-Gal4* was used for *baboA-*KD in the medulla precursors. Anti-24B10 labels axonal track of photoreceptor neurons (R1-R8) in the medulla field in *w^1118^* (con) and *baboA-*KD. (A,B) Anti-24B10 patterns in the brain of con (n=7) and *baboA*-KD (n=7) at WL3 stage. The labeled Volwig’s nerve (yellow arrowhead in B) is derived from the transgenic background of the *scro-Gal4* line. Aberrant targeting is indicated by a white arrowhead. (C-Ciii) Normal anti-24B10 patterns in the adult optic lobe shown from optical z-sections (n=7). (Ciii) Co-labeling of Anti-24B10 (red) and DAPI (white) of Cii. (D-Diii) Optical Z-sections showing severely defective anti-24B10 patterns in the *baboA*-KD (n=7). (Diii) Anti-24B10 staining with DAPI stains in Dii. Empty spaces (*) are formed by severe mis-routing of R7-R8 axons. (E-Eii) mCD8GFP expression merged with anti-24B10 staining shown in (D-Dii) sections. (Eiii) mCD8GFP only of Eii. Scale bars in A and C, 50 μm.

### Characterization of *babo^Gal4^* driver

Gal4 drivers are essential and versatile for various transgenic manipulations of gene expression. Thus far, there is no *babo*-specific Gal4 driver. A *babo^CR00274-TG4.1^* line (shortly, *babo^Gal4^*) has a T2A-Gal4 Trojan exon inserted in the second and common intron (Fig 16A) [63]. To test if this line can serve as *babo*-specific Gal4 driver, we studied reporter gene expression patterns driven by this line. Expression of nuclear GFP (nGFP) is found in a wide range of cell types including NEs and NBs and their lineage cells as well as MB neurons and larval neural cells with heterogenous expression levels (Fig 16B-16E), and the overall patterns are comparable to pan-Babo-GFP ones. Differential reporter levels in various cell types derived from postembryonic neurogenesis are also reminiscent of the BaboA-GFP ones (Fig 16B-Bii, 16C-Cii). However, we also noted an incongruity in that *babo^Gal4^-*reported expression lacks in some of CB-NBs and their lineage cells as well as in some of OPC (Fig 16C-Cii, 16D, 16E, 16F-Fi). To determine if this is due to the UAS-nGFP line that we employed, we tested a UAS-redstinger (nRFP) line, which, in our hands, shows stronger signals and found that some of CB-NBs and OL-NEs are still devoid of nRFP expression (Fig 16E, 16F-Fi, 16G-Gii). As expected, *babo^Gal4^* activity is found in the MB neurons (Fig 16H-Hii) and in large population of glial cells both on the surface and internal CNS regions (S4 Fig). However, *babo*>reporter expression is not observed in some of Pros-positive ag glia of the VNC (S4 Fig). We reasoned that certain incongruities might be explained by difference in the half-life of Babo proteins vs. reporter proteins or cell type-specific alteration in transcriptional activity or splicing pattern.

**Fig 16.**
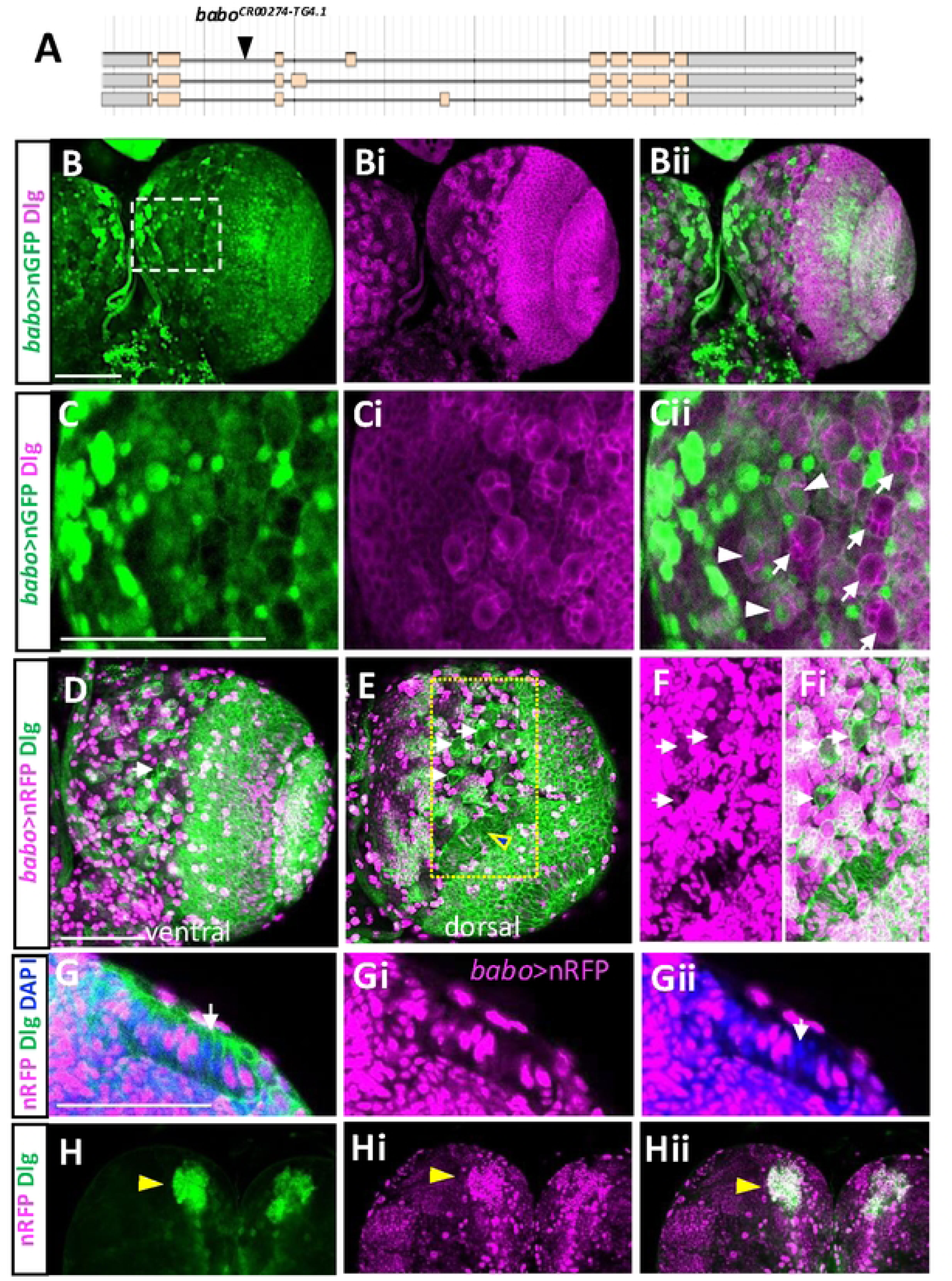
Reporter expression driven by *babo^Gal4^* in the larval brain. (A) Insertion site of a *T2A-Gal4* construct in the 2^nd^ intron of *babo*. (B-Bii) Double-labeling of *babo*>nGFP (green) and Dlg (magenta) is shown in the ventral side of a brain lobe (n=8). (C-Cii) Magnified images of a boxed region in (B). Arrows indicate *babo*>nGFP signals lacking in NBs and their lineage cells. Arrowheads indicate NBs and their clones containing *babo*>nGFP expression. (D,E) A brain lobe showing *babo>*nRFP and Dlg from the ventral (C) and dorsal (D) sides (n=8). Composite of three dorsal z-stacks (each about 2 μm thick). Arrows indicate NBs lacking *babo>*nRFP expression. An arrowhead indicates NEs lacking *babo>*nRFP. (F-Fi) Magnified images in a boxed region in (E) with longer exposure to detect low levels of *babo*>nRFP expression. (G-Gii) Some NEs labeled with Dlg and DAPI lack *babo>*nRFP expression. (H-Hii) *babo*>nRFP in the Dlg-labeled MB neurons. Scale bars, 50 μm.

### Knockdown phenotypes of individual isoforms by *babo^Gal4^*

We also examined mutant phenotypes of *babo^Gal4^*. The *babo*-null mutants mostly die during the larval stages, but rare escapers manage to develop to small tubby-like prepupae that die during the early pupal stage (Fig 2A). In contrast, a small population of homozygous *babo^Gal4^* mutants develop up to the pharate stage (Fig 17A vs. 17B). Their overall pupal phenotypes, such as elongated body, short appendages, cuticle hardening defect, and eclosion failure, resemble *baboA-indel* and *babo>baboA-miRNA* flies rather than *babo*-null mutants (Fig 17B-17D) [10]. These observations suggest that *babo^Gal4^* is a hypomorph allele.

**Fig 17.**
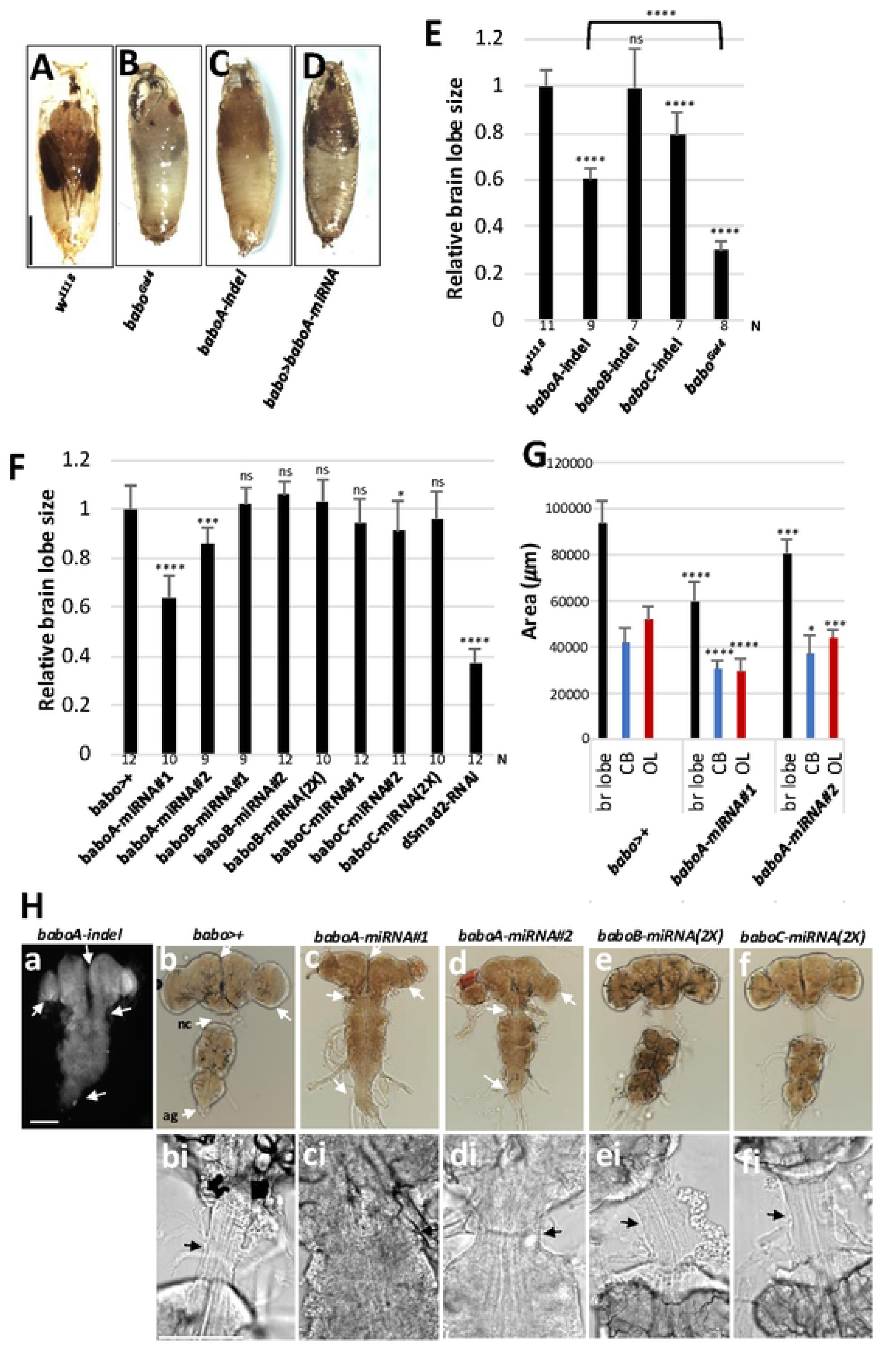
*babo^Gal4^* is a hypomorphic allele and a potent *babo-gal4* driver. (A-D) Pharate pupal phenotypes of *w^1118^* (con), *baboA-indel* (n=10), *babo^Gal4^* (n=5), and *babo>baboA-miRNA* (n=43). (E) A graph showing relative brain lobe sizes of *w^1118^*, individual isoform mutants, and *babo^Gal4^* homozygous mutants at 120 h AEL. Numbers of samples (N) for each genotype are shown under the bars. (F) A graph showing relative brain lobe size of *babo^Gal4^* crossed to *w^1118^* (*babo*>+, con), two different *baboA-miRNA* lines (#1, #2), two copies (2X) of *baboB-* and *baboC-miRNA*, and *dSmad2-RNAi* at 120 h AEL. More than six samples for each genotype were used for quantification. (G) Areas of brain (br) lobe, central brain (CB), and optic lobe (OL) measured on ventral to dorsal middle sections. (H) Pharate CNSs of indicated genotypes. (a-f) White arrows point to defective fusion of the two brain hemispheres, separation of the subesophageal ganglia from the thoracic ganglion and formation of the neck connective (nc), growth of optic lobe, and condensation of abdominal ganglia (ag) in *A-indel* mutants and *baboA*-KD. (bi-fi) Magnified images of the neck connective area (arrow) shown in (b-f). Scale bars: in B, 1 mm; in Ha, 100 μm; in Hbi, 50 μm.

We further compared the small brain phenotypes of this allele with the isoform mutants. Loss of pan-Babo function severely affects postembryonic neurogenesis, resulting in the most severe small brain phenotype [10, 27, 36]. *baboA-indel* also impairs larval brain growth, although the small brain phenotype of *baboA-indel* mutants is less severe than that of *babo*-null [10]. *baboC-indel* mutants display slightly reduced brain size at 120 h AEL (Fig 17E); this is mainly due to their delayed larval development [10]. Overall, a small brain phenotype of *babo^Gal4^* is significantly more severe than those of *baboA-indel* mutants (Fig 17E).

Next, we examined the brain sizes of larvae carrying *babo^Gal4^*-driven KD of each *babo* isoform at 120 h AEL. Two different *miRNA* lines were tested for each isoform. The *baboB*-KD, even with two *miRNA* copies, has no significant impact on the brain size (Fig 17E and 17F), which is consistent with *baboB-indel* mutants [10] and the lack of expression in the CNS. The *baboC*-KD, like *baboC*-indel, shows only slightly smaller brains, regardless of *miRNA* dosage (Fig 17E and Fig 17F). Both *baboA-miRNA* lines display developmental defects, such as the lack of wandering behavior, pupariation on food surface, defective puparia formation (tanning and constriction of pupal body), and pupal lethality, although #1 has slightly more severe defects than #2 in certain aspects of brain development (Fig 17F and 17G). Overall *baboA*-KD phenotypes are comparable to those of *baboA-indel* mutants [10].

We discovered an additional *baboA*-specific CNS phenotype. *baboA-indel* mutants that developed to the pharate stage exhibit noticeable defects in metamorphosis-related CNS reorganization such as separation of the subesophageal ganglion (SEG) from the thoracic ganglion (TG) with concomitant formation of the neck connective (a.k.a. cervical connective), fusion of the brain lobes, and condensation of the abdominal ganglia (Fig 17Ha vs. 17Hb). The *baboA*-*mRNA* #1 mimicks the phenotypes of *baboA*-*indel* mutants more closely, while *baboA-mRNA* #2 shows partial defects (Fig 17Hb-d, bi-di); such differential effects by the two *miRNAs* are consistent with larval brain growth defect. Neither *baboB*-KD nor *baboC*-KD has any of these defects (Fig 17He-f, ei-fi). Considering the effectiveness of both *baboB-miRNA* and *baboC-miRNA* (Fig 4) and CNS-expression of the three isoforms, we conclude that BaboA is the most influential isoform for most aspects of metamorphosis-related CNS organization.

## Discussion

Various *babo* mutations produce multifaceted defects in developmental, physiological, behavioral, and morphological processes, suggesting that the Babo-mediated TGF-β signaling plays diverse roles in different tissues including the CNS during larval development and metamorphosis [8, 10, 36, 64, 65]. In addition to these genetic studies, investigating the spatio-temporal expression of individual Babo isoforms is crucial to understand their functions in various developmental contexts. For this aim, we generated the GFP-tagged transgenic lines to probe endogenous production of the isoforms. In this study, we demonstrate that the three Babo isoforms are differentially expressed in both time (life stage) and space (tissue- and cell-type specificity), with a focus on the RG and CNS that are important for the timing of insect development and behavior. Beyond the tissues examined here, Activin signaling functions in other developmental context, and uncovering the isoform expression patterns in combination with cell-specific knockdown assays will facilitate functional studies. Our findings highlight that these reporter lines, along with *babo^Gal4^*, are valuable tools that can help delineate various functions of the isoforms in contributing to the different aspects of the *babo* phenotype.

### Subcellular targeting of GFP-conjugated Babo proteins

All endocrine tissues of the larval RG display overlapping expression of both BaboA and BaboC, but the two exhibit clear differences in their expression levels depending on endocrine type. Epidermal cells also express both BaboA and C with BaboC being more predominant. Interestingly, the larval muscle cells express all isoforms in the order of BaboB> BaboA> BaboC. Our observations signify sophisticated molecular strategies operating in a tissue- (or cell-) and a stage-specific manner at the splicing event to specify not only isoform types but also differential levels. Therefore, it would be interesting to explore molecular mechanisms as to how the splicing event is biased/regulated to determine single or multiple isoforms [66, 67].

We found that the isoform-specific GFP tagging influences the subcellular trafficking. The C-terminally tagged isoforms (pan-Babo-GFP) appear properly targeted to the plasma membrane in all tissue types we examined indicating that the C-terminal tag does not disrupt the trafficking process. However, BaboA-GFP is found in the cytoplasm, which would negate its function as a receptor. On the other hand, BaboC-GFP targeting varies depending on cell type; it is found in the cytoplasm of neurons and other tissues, but in the processes of glial cells. Similarly, BaboB-GFP is cytoplasmic in the adult CA cells but on the membrane of some gut cells. One exception is the muscle cells that all tagged isoforms found at the NMJ. Although the nature of the proper targeting of the tagged isoforms in the musclular cells is not clear, we speculate them to be functional as receptors. These observations show that internal GFP tag does affect subcellular targeting differently based on cell type and isoform type. It also explains why pan-Babo-GFP rescued the lethality of *babo*-null mutants but the rescue activity of isoform-GFP lines depends on the tagged isoform.

### BaboC in the CNS

Genetic studies have shown that the Daw/BaboC-mediated TGF-β signaling regulates circulating sugar concentration, total triacylglycerol, and glycogen content [8] and that loss of this isoform function slows down larval development [10]. In this study, we demonstrate that BaboC is expressed in a large population of larval neurons in the suboesophageal, thoracic and abdominal ganglia, where motoneurons are densely populated. The suboesophageal ganglion (SEG) not only harbors motoneurons but also receives and processes gustatory information, all of which are important for regulating larval feeding behavior [68–70]. Furthermore, BaboC is also expressed in the insulin-producing cells as well as in the CC which is the only source for AKH [32, 37]. These results altogether support the idea that the Daw/BaboC signaling is involved in regulation of energy metabolism or/and feeding behavior to promote proper larval growth.

Another major site of BaboC expression is larval glia. At least three glial subtypes are tied with its expression: pg, cg, and ag. The pg play a role in forming the BBB together with another surface glia subtype, the subperineurial glia (spg), sensing body-generated circulating signals, and importing necessary nutrients, metabolites and ecdysone hormone [39, 71–75]. The pg is also the only source for Daw in the brain [11, 13]; hence, it is plausible to propose that BaboC function in the pg might be involved in modulation of brain Daw levels in an autocrine manner in response to the circulating ones. Another BaboC expression site, the cg, is known to provide necessary cues that influence activities of NBs and NEs [76–78]. Therefore, this isoform could play a certain role in functional and/or developmental aspects of postembryonic neurogenesis in coordination with the developmental status of the whole body. In line with this, a loss of BaboC function delays brain growth in parallel to delayed larval development [10].

### Roles of BaboA for adult CNS development

BaboA is widely expressed in larval neurons, and such patterns fit well with its broad roles reported in larval brain development [10]. Two major metamorphosis-related cellular events in larval neurons have been ascribed to BaboA function. The MB γ neurons prune their axons and dendrites during early phase of metamorphosis in order to reform adult-specific ones, and the Myo/BaboA-mediated TGF-β signaling is essential for such remodeling process [7, 64]. In this study, we revealed that the same signaling is also required for remodeling of peptidergic larval neurons. In *baboA-indel* mutants these neurons fail to shed larval patterns of the neural circuit, thus unable to form adult patterns. Based on these data, we propose that BaboA-mediated neuronal remodeling is not specific to the MB γ neurons but rather general to other persisting larval neurons. This signaling also plays a key role in triggering PCD of vCrz neurons that occurs soon after puparium formation [9, 79]. The rescued vCrz neurons by expression of P35-a universal inhibitor of caspases are also subjected to remodeling, indicating that it is the default process for all of persisting larval neurons [49, 50]. In contrast, the undead vCrz neurons in *baboA-indel* mutants maintain juvenile neural circuitry, further supporting a general role for BaboA in the remodeling process of persisting larval neurons. Another question is, then, whether BaboA function is also a general requirement for the removal of the obsolete larval neurons by developmental PCD. A large population of motoneurons and interneurons in the larval VNC undergo apoptosis during the early phase of metamorphosis since their functions are no longer needed in the adult stage due to changes in locomotion style (from crawling to walking and flying), which results in a dramatic shrinkage of the abdominal ganglia [80–82]. The abdominal ganglia of *baboA-indel* pharates are substantially reduced in volume, but it is not as severe as in wild-type, arguing that many obsolete larval neurons still undergo PCD even in the absence of BaboA function. Therefore, it is unlikely that BaboA-mediated TGF-β signaling is a general requirement for PCD of larval neurons that are programmed to die by pharate stage.

Prominent expression of BaboA is also found in all cell types of postembryonic neurogenesis. The loss of this isoform function was shown to reduce the proliferation rate of both NBs and NEs as well as NE-to-NB conversion, two events which lead to generation of a proper population of new neurons necessary to build an adult brain, suggesting it plays a role in both cellular events [10]. Together with previous stage-and cell-specific KD assays and this study showing expression of BaboA in both types of neural progenitors strongly support the notion that this isoform acts autonomously for the events. Once precursor neurons are generated, they must complete their differentiation by establishing highly interconnected neural circuits in the adult brain. BaboA is detected in all precursors from their birth, suggesting that this isoform may play roles in the process of neuronal identity establishment and differentiation. Indeed, BaboA function has been shown to establish the proper neural circuitry of dorsal cluster neurons in a postembryonic NB lineage labeled by *atonal-gal4* [65]. We also found that loss-of-function *baboA* mutants are devoid of adult-specific NPFergic neuronal clusters, but not those of CRZ, DSK and PDF. Instead, BaboA appears to function to maintain high levels of PDF expression in adult-specific PDF neurons. These results lead us to conclude that BaboA function is involved in diverse postembryonic neurogenesis events and functions of adult neurons in a cell-specific manner.

BaboA is also important for the cell-type specification or differentiation of the medulla precursor neurons. In the adult optic lobe, anti-24B10 marks axons of the retinal R7 and R8 that terminate at distinct medulla layers, M6 and M3, respectively [62]. The evenly spaced trajectories of the retinal axons to their M3 and M6 layers also mirror columnar medulla neuropil organization by the medulla neurons [83]. Our results suggest that *baboA*-depleted medulla neurons fail to establish the M3 and M6 layers, unique features of the adult medulla neuropil. This indicates that BaboA function is required for proper development of the medulla neurons, either for cell-type specification or connectivity or both. Alternatively, the BaboA-mediated signal might be a key player in coordinated differentiation of the medulla neurons during metamorphosis in response to the arrival of new afferent input [84]. For instance, the medulla neurons might undergo two-step differentiation processes as shown by the pioneer neurons of four type II NB lineages to establish the central complex neuropil structures, juvenile one (primary) and then adult one (secondary) [85, 86]. If BaboA-mediated signaling provides an essential cue for the later coordinated step, we expect that medulla neurons would be unable to elaborate adult patterns of medulla neuropil in the absence of BaboA, thus maintaining primary ones. This also explains the absence of layer formation in the adult optic lobe of *scro>baboA-miRNA* flies and the uncanny resemblance of the defective patterns to larval ones.

Underlying anatomical and behavioral differences observed between larval and adult stages in holometabolous insects are alterations in CNS structure and function [87]. Several large-scale morphogenetic features of such CNS metamorphosis include: SEG-TG separation along with formation of a thin and long cervical connective, condensation of abdominal ganglia due to massive PCD of abdominal larval neurons, fusion of the brain lobes at the midline, and elaboration of highly ordered four neuropils of the OL in combination with size increase [84, 88–90]. Of interest, both *baboA-indel* mutants and *babo>baboA*-KD flies exhibit defects in these morphogenetic reorganizations. One way to interpret these data is that BaboA-mediated TGF-β signaling regulates stage-specific upregulation of both EcR-B isoforms [7, 10, 64]. In conclusion, our study supports previously suggested roles of BaboA and uncovers a broad range and novel roles for BaboA in re-construction of a larval CNS to an adult one involving neurons of both embryonic and postembryonic origin. The reagents described here should also help parse out the specific functions of individual isoforms in many other tissues and cell types during development.

## Acknowledgments

We greatly appreciate Dr. R. Goodchild for the kind gift of rabbit-anti-GFP, Dr. P. Leopold for rat-anti-Dilp2, and Dr. Z. Lee for UAS-babo-miRNA plasmids and fly lines.

## Supporting information

**Table S1.** Pairs of primers to produce C-terminal and isoform-specific Babo-EGFP conjugation. *babo* tails in the primers are shown in red, and plasmid sequences are in black--*C term PL452 for* and *C EGFP* for C-terminal construct; *N term PL452 for* and *N EGFP rev* for isoform-specific constructs.

**S1 Fig**. **GFP signals in the larval CNS and RG from homozygous *babo^fTRG00444.sfGFP-TVPTBF^* flies.** WL3 larvae homozygous for the *babo^fTRG00444.sfGFP-TVPTBF^* construct were processed with anti-GFP and anti-Dlg for the CNS (n=14) and anti-GFP alone for the RG (n=7). (A-C) Ventral side of the brain lobes showing anti-GFP (A), anti-Dlg (B) signals, and merged one (C). (D) Anti-GFP immunostaining (red) in the larval RG. Membrane-bound Babo-fusion proteins are detected in all cell types of the RG. Scale bars in A, 100 μm, and in D, 50 μm.

**S2 Fig**. **Expression of Pan-Babo-GFP and GFP-tagged isoforms in the adult CNS.** GFP staining in the anterior side of the adult brain (A-E) and ventral side of the VNC (Ai-Ei) of indicated genotypes. The whole CNSs were dissected from one-two week-old adult flies homozygous for each transgene, processed for anti-GFP labeling, and imaged simultaneously under the same condition (n>7, for each genotype). Scale bars in A and Ai, 50 μm.

**S3 Fig**. **Detection of BaboA-GFP in the developing larval CNS.** (A-E) BaboA-GFP at 24 h, 48 h, 72 h, 96 h and 120 h after egg laying (AEL). Scale bars in A, E, Ai, Bi, and Ci, 50 μm. (Ai-Ci, Ei) Enlarged images showing double-labeling of BaboA-GFP and anti-Dlg. Scale bars in A, E, Ai, Bi, and Ci, 50 μm.

**S4 Fig. Reporter expression driven by *babo^Gal4^* in glial cells.** (A,B) Expression of mCD8GFP driven by *babo^Gal4^* in surface (A) and cortex glial cells (white arrows, B). (C-Cii) Surface glia on the dorsal-most surface of a brain lobe showing co-expression of Repo and *babo>*nRFP. Nearly all surface glial cells show nRFP signals weakly or strongly. (D-Dii) Some Repo-positive glial cells devoid of *babo>*nRFP in dorsal side of the VNC. (E-Eii) *babo>*nRFP expression is not detected in some of Pros-positive astrocyte-like glial cells. Scale bars, 50 μm.

